# Inferring MHC interacting SARS-CoV-2 epitopes recognized by TCRs towards designing T cell-based vaccines

**DOI:** 10.1101/2020.09.12.294413

**Authors:** Amir Hossein Mohseni, Sedigheh Taghinezhad-S, Bing Su, Feng Wang

## Abstract

The coronavirus disease 2019 (COVID-19) is triggered by severe acute respiratory syndrome mediated by coronavirus 2 (SARS-CoV-2) infection and was declared by WHO as a major international public health concern. While worldwide efforts are being advanced towards vaccine development, the structural modeling of TCR-pMHC (T Cell Receptor-peptide-bound Major Histocompatibility Complex) regarding SARS-CoV-2 epitopes and the design of effective T cell vaccine based on these antigens are still unresolved. Here, we present both pMHC and TCR-pMHC interfaces to infer peptide epitopes of the SARS-CoV-2 proteins. Accordingly, significant TCR-pMHC templates (Z-value cutoff > 4) along with interatomic interactions within the SARS-CoV-2-derived hit peptides were clarified. Also, we applied the structural analysis of the hit peptides from different coronaviruses to highlight a feature of evolution in SARS-CoV-2, SARS-CoV, bat-CoV, and MERS-CoV. Peptide-protein flexible docking between each of the hit peptides and their corresponding MHC molecules were performed, and a multi-hit peptides vaccine against the S and N glycoprotein of SARS-CoV-2 was designed. Filtering pipelines including antigenicity, and also physiochemical properties of designed vaccine were then evaluated by different immunoinformatics tools. Finally, vaccine-structure modeling and immune simulation of the desired vaccine were performed aiming to create robust T cell immune responses. We anticipate that our design based on the T cell antigen epitopes and the frame of the immunoinformatics analysis could serve as valuable supports for the development of COVID-19 vaccine.

## Introduction

The severe acute respiratory syndrome coronavirus 2 (SARS-CoV-2) is determined as the causative agent of coronavirus disease 2019 (COVID-19) by public health professionals [1]. At the time of writing this manuscript, researchers tend to focus their attention on describing the immune response against SARS-CoV-2-derived B and T cell epitopes [2–4]. Although considerable efforts have been spent on clarifying the SARS-CoV-2 epitopes, the used approaches often lack accurate T cell receptor (TCR) and peptide-bound major histocompatibility complex (pMHC) binding models, which are crucial to initiate adaptive immunity [5–10]. Thus, little is known about the relationship between an effective adaptive immune response and the SARS-CoV-2 epitopes.

The recognition of antigen is a critical factor in T cell activation [11]. It has been well documented that the TCR plays a key role in activating cell-mediated adaptive immune responses. In this process, the TCR recognizes a peptide, obtained from a protein antigen, presented by MHC molecules, resulting in the initiation of downstream signaling pathways within the T cell. Moreover, immunological studies have found that MHC molecules will present selective peptides derived from a protein, indicating the significant impact of predicting peptide-MHC binding towards identification of potential T cell epitopes [12]. These findings acknowledge the necessity of further studies to provide insights into TCR–pMHC interactions and binding mechanisms in order to select optimal peptide antigen (Hit peptide) for eliciting potent immune responses against infectious diseases [13, 14]. In keeping with these observations, deep understanding of the peptides originating from the SARS-CoV-2 proteins that could elicit T cell mediated immune response may help in quest for the better development of a rational design of COVID-19 vaccine. To address these issues, we applied a novel computational framework to calculate the interaction between TCRs and SARS-CoV-2 antigen peptides whose recognition is restricted by human leukocyte antigen (HLA). Efficient structural modeling of TCR-pMHC interaction from SARS-CoV-2 proteins could identify the appropriate hit peptide(s), that with the following immunoinformatics analysis relating to the potential epitopes, is expected to provide a beneficial data source for the development of COVID-19 vaccine.

## Materials and Methods

### Retrieving protein sequences

All full-length protein sequences of envelope protein (E), membrane glycoprotein (M), nucleocapsid phosphoprotein (N), open reading frame 1ab polyprotein (ORF1ab), ORF3a protein (ORF3a), ORF6 protein (ORF6), ORF7a protein (ORF7a), ORF8 protein (ORF8), ORF10 protein (ORF10), and surface glycoprotein (S) of SARS-CoV-2 in FASTA format from different geographic regions including: China, Wuhan (GenBank: MN90894) (as a reference SARS-CoV-2 isolate), India (Genbank: MT050493), Italy (Genbank: MT066156), Nepal (Genbank: MT072688), Sweden (Genbank: MT093571), Brazil (Genbank: MT126808), China (Genbank: MT135041), Taiwan (Genbank: MT192759), Viet Nam (Genbank: MT192773), Spain (Genbank: MT198652), Pakistan (Genbank: MT240479), Colombia (Genbank: MT256924), Peru (Genbank: MT263074), USA (Genbank: MT276329), and South Korea (Genbank: MT304474) were retrieved from the GenBank (https://www.ncbi.nlm.nih.gov/genbank/) at the first of April 2020. Also, the entire viral proteome sequences from bat-CoV (Genbank: MG772921), MERS-CoV (Genbank: KF958702), and SARS-CoV (Genbank: AY390556) were retrieved for further comparison between the SARS-CoV-2 derived hit peptides and other coronaviruses.

### Modeling of TCR–pMHC complex

The sequences of each protein were aligned using the CLC sequence viewer (Version 6.7.1). For computing the final alignment, the sequences showing poor alignments were deleted and finally the consensus sequences of each protein were extracted for using as an input for modeling of TCR–pMHC. Then, each protein sequence in FASTA format was allocated by PAComplex server to forecast a model of 9- to 11-mer peptides bound to HLA-A0201, HLA-B0801, HLA-B3501, HLA-B3508, HLA-B4405, and HLA-E-peptide-TCR template with Z values > 4.0 [15]. Moreover, an in-depth interaction among atoms, model of binding, and amino acid sequences related to homologous peptide antigens were evaluated for each query. Then, in order to evaluate the conserved regions and pairwise identity among SARS-CoV-2-derived hit peptide and other coronaviruses including SARS-CoV, MERS-CoV, and bat-CoV multiple sequence alignment (MSA) was performed.

### Prediction of immunogenicity of the SARS-CoV-2-derived hit peptide

Each SARS-CoV-2-derived hit peptide by same length that presented on the same HLA class I molecule were analyzed through evaluation of the amino acid characteristics by T cell class I pMHC immunogenicity tool [16]. Prediction results were sorted by descending score values. The higher score was indicated as a greater probability of eliciting an immune response.

### Study the binding mechanisms and interatomic interactions

For each hit peptides, hydrogen bonds (H-bond) and van der Waals (VDW) forces of pMHC and peptide-TCR interfaces were calculated. Also, in order to study the atom–atom contacts, Arpeggio server was used for detailed evaluation of TCR-pMHC complex derived from SARS-CoV-2 proteins. Accordingly, protein structure and visualization of the measured interactions between atoms including the strongest mutually exclusive interactions, polar contacts, H-bonds, ionic interactions, aromatic contacts, hydrophobic contacts, and carbonyl interactions were showed by WebGL-based protein structure viewer and PyMOL session based-visualization.

### Peptide-protein flexible docking

One of the significant tools for designing of drug is computational docking methods. So, the usage of innovative methods including protein-peptide docking for quick development of peptide therapeutics in rational drug design is unavoidable. In this study, template-based docking and 3D protein-peptide complex structures from input protein structure and peptide sequence were predicted by GalaxyPepDock server (http://galaxy.seoklab.org/cgi-bin/submit.cgi?type=PEPDOCK) through merging information on analogous interactions in the structure database and energy-based optimization in order to expect the formation of MHC-peptide complex. Then, the contacting residues in the template structure to the template amino acids was aligned by similarity of the amino acids of the target complex and an interaction similarity score was defined for it by server. This analysis presented an example of MHC-peptide docking performed by each individual hit peptides derived from N and S proteins and available PDB file of HLA alleles including HLA-A0201 (PDB: 5yxn), HLA-B3501 (PDB: 1a9e), HLA-B0801 (3spv), HLA-B3508 (3bw9), HLA-B4405 (3dx8), and HLA-E (2esv), separately.

### Designing of multi-hit peptides vaccine sequence

A set of high immunogenic hit peptides derived from N and S proteins of SARS-CoV-2 with high binding events to HLA-A0201, HLA-B0801, HLA-B3501, HLA-B3508, HLA-B4405, and HLA-E were selected on the basis of their solvent exposed residues and hydrophobicity scales. The AAY and GPGPG linkers were applied for linking the candidate N and S hit peptides together, respectively. Additionally, the human beta defensin 3 was also joined at N-terminus of the vaccine construct using EAAAK linker which acts as adjuvant to improve the immunogenicity of the multi-hit peptides vaccine.

### Prediction of allergenicity, antigenicity, and physiochemical properties of designed vaccine

The allergenicity of query sequence was evaluated using AllergenFP v.1.0 server (http://ddg-pharmfac.net/AllergenFP/). Vaccine construct was analyzed by VaxiJen v2.0 (http://www.ddg-pharmfac.net/vaxijen/VaxiJen/VaxiJen.html) with high accuracy at 0.4 thresholds for virus. Also, ProtParam web server (https://web.expasy.org/protparam/) was used to predict various physicochemical properties of vaccine construct including like theoretical isoelectric point (pI), in vitro and in vivo half-life, amino acid composition, molecular weight, instability and aliphatic index, and grand average of hydropathicity (GRAVY).

### Vaccine-structure modeling

The SOPMA web server (https://npsa-prabi.ibcp.fr/cgi-bin/npsa_automat.pl?page=npsa_sopma.html) was utilized for the computation the secondary structure of multi-hit peptides vaccine. Indeed, I-TASSER server (https://zhanglab.ccmb.med.umich.edu/I-TASSER/), as an algorithm for fast and accurate de novo protein structure prediction, was further employed to predict the solvent accessibility, 3-dimensional structure (3D), and function of the vaccine sequence. In detail, I-TASSER by recruiting SPICKER program groups all the decoys according to the pair-wise structure similarity and provides five models that are linked to the five largest structure clusters. This server determines the confidence of each model by C-score which typically is in the range of (−5, 2). A model with a higher confidence has higher C-score than a model with a lower confidence. Also, prediction of TM-score and RMSD was done by I-TASSER server according to the C-score and protein length after obvious connection between these qualities.

### Refinement of tertiary structure and validation

The refinement of the tertiary structure of vaccine was carried out using 3D^refine^ server (http://sysbio.rnet.missouri.edu/3Drefine/index.html) which is based on the CASP10 tested refinement method for protein's side chain reconstruction, molecular dynamics simulation, and repacking to relax the 3D structure. Then, to fulfill the validation of 3D structure, output model of 3D^refine^ server was subjected to the ProSA-web server (https://prosa.services.came.sbg.ac.at/prosa.php). Overall quality of the model with Z-score index was anticipated for model by the server. The outside range of Z-score value for the predicted model will point to the invalid structure of the model. Indeed, RAMPAGE server (http://molprobity.biochem.duke.edu/) also was used to determine the overall quality of the predicted model of the multi-hit peptides vaccine following Ramachandran plot analysis.

### Immune simulation

To further characterize immune response profile of multiple hit-peptide protein, FASTA format of our multi-hit peptides vaccine was subjected to C-ImmSim server (http://150.146.2.1/C-IMMSIM/index.php) for computing T cell immune response. C-ImmSim is an agent-based model that can predict immune epitope and immune interactions by using PSSM (a position-specific scoring matrix) and machine learning techniques, respectively. It instantaneously induces three compartments in three distinct anatomical regions (bone marrow thymus, a tertiary lymphatic organ). As such, three injections with time steps set at 1, 84, and 168 were given four weeks apart along an injection without LPS. Also, host HLA were selected for HLA-A0201, HLA-A0101, HLA-B0801, HLA-B3508, and HLA-DRB1_0101. Nevertheless, the simulation volume and simulation steps were set based on the default parameters.

### Statistical analysis and data availability

In our theoretical study, no statistical analyses were performed according to the literature data and web-accessible databases. All data presented in the present study are summarized in the complementary figures, tables, and supplemental materials.

## Results and Discussion

### Overview of the potential TCR–pMHC binding models for SARS-CoV-2 proteins

According to TCR–pMHC binding models, for N query, one hit peptide antigen candidate (105SPRWYFYYL113) with Z-value cutoff > 4 and 61 homologous peptide antigens in 34 organisms was found by using HLA-A0201-peptide-TCR template [PDB entry 2vlr] and the experimental peptide database. It was documented that E, M, and ORF10 protein queries have two hit peptide antigen candidates (Z-value cutoff > 4). Also, 108, 80, and 40 homologous peptide antigens in 56, 40, and 25 organisms by using HLA-A0201-peptide-TCR template [PDB entry 1qrn, 2jcc, and 1oga] and the experimental peptide database was identified for them, respectively. Giving our data, five and ten hit peptide antigen candidates (Z-value cutoff > 4), and 23 and 40 homologous peptide antigens in 16 and 25 organisms by using HLA-A0201-peptide-TCR template [PDB entry 2p5e and 1oga] and the experimental peptide database were inferred in ORF3a and S proteins, respectively. No hit peptide antigen candidates with Z-value cutoff > 4 were detected for ORF6, ORF7a, and ORF8 queries by using HLA-A0201-peptide-TCR template (Figure S1A). Our results showed that by using HLA-B0801 (Figure S1D), HLA-B3501 (Figure S1B), and HLA-B3508 (Figure S1E) template, TCR-pMHC models with Z-value cutoff > 4 were only predicted for S (5 hit peptide antigen candidate and 91 homologous peptide antigens in 57 organisms with PDB entry 1mi5), N (3 hit peptide antigen candidate and 3 homologous peptide antigens in 2 organisms with PDB entry 2nx5), and S (1 hit peptide antigen candidate and 1 homologous peptide antigens in 1 organisms with PDB entry 2ak4) proteins, respectively. While, only M and S proteins were used to predict TCR-pMHC complex by using HLA-B4405 (Figure S1C) with PDB entry 3dxa (15 homologous peptide antigens in 12 organisms). Also only N, ORF10, and S proteins were used for prediction of TCR-pMHC complex by HLA-E (Figure S1F) with PDB entry 2esv (84 homologous peptide antigens in 41 organisms) with Z-value cutoff > 4. No hit peptide antigen candidates with Z-value cutoff > 4 were detected for M, E, and ORF3a queries by using HLA-E-peptide-TCR template (Figure S1F). Moreover, model for ORF1ab query did not predict due to its sequence length was ≥ 300. The 3D structure of the TCR–pMHC complex for each hit peptide derived from each protein was illustrated by SWISS-MODEL. Detailed information about derived hit peptides for SARS-CoV-2 along with bat-CoV, MERS-CoV, and SARS-CoV were tabulated in Table 1.

**Table 1:**
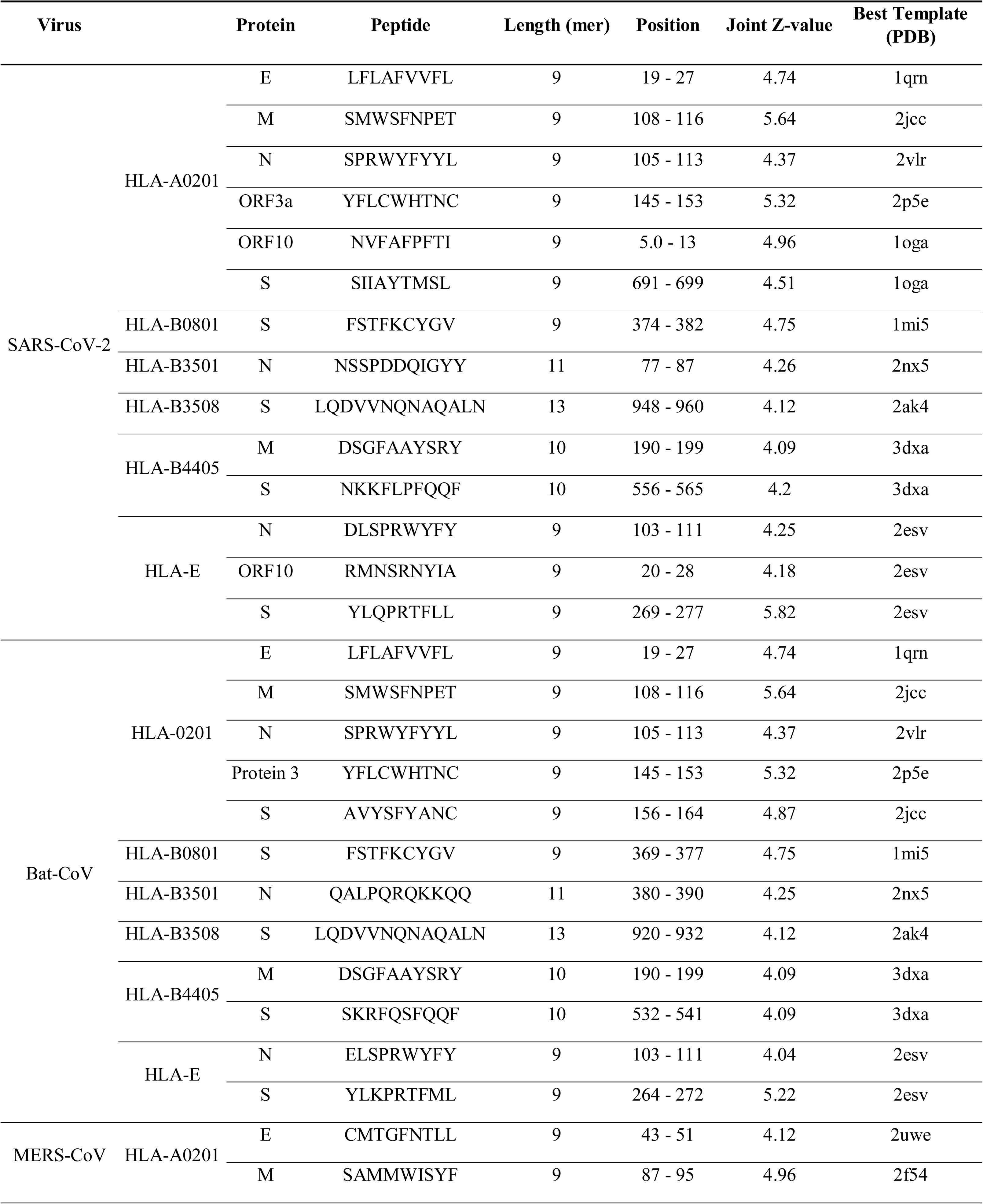
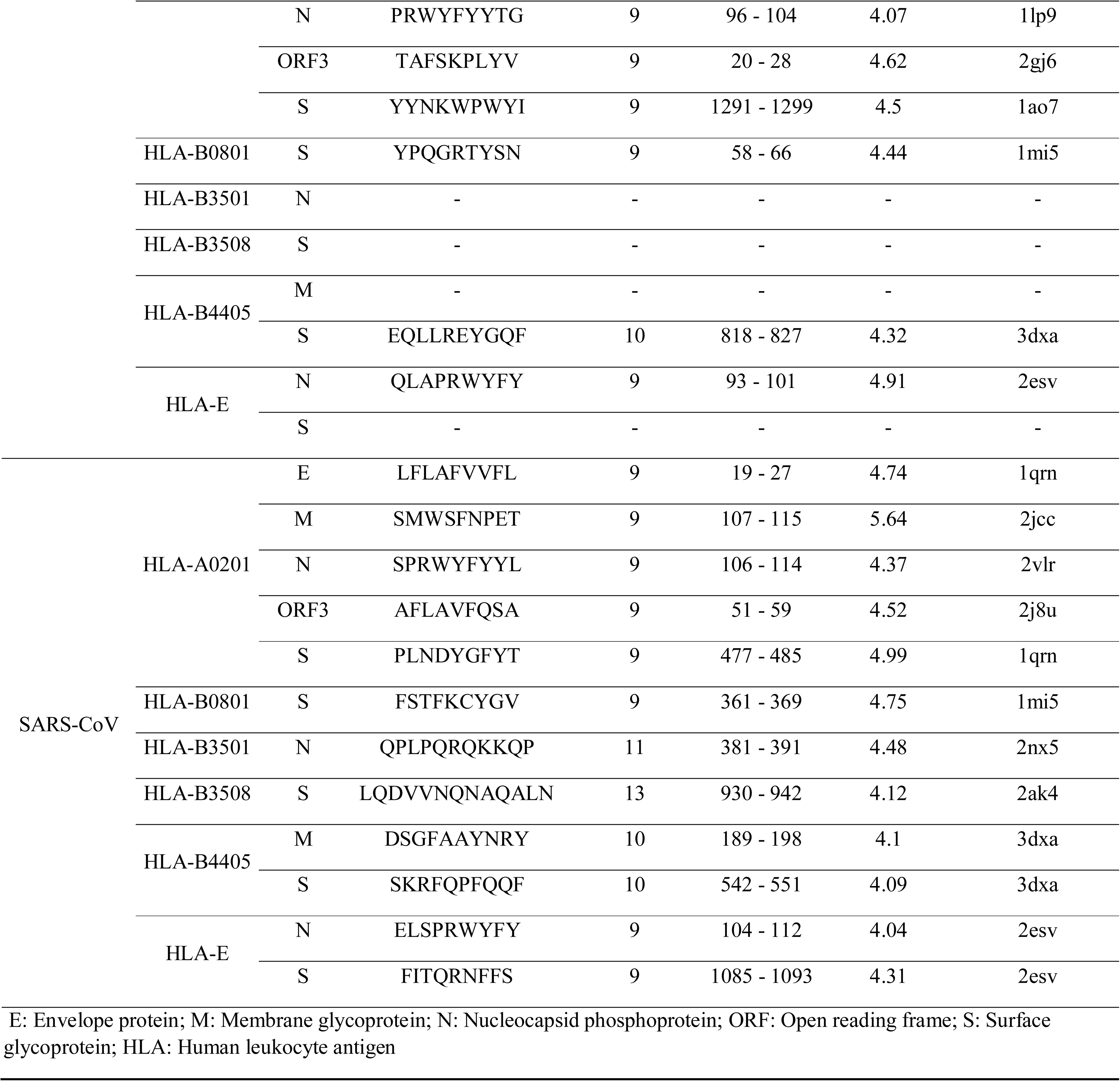
Hit peptide derived from TCR-pMHC complex.

### The binding events among hit peptides derived from N proteins related to HLA-A0201

Our TCR-pMHC models predicted that position 2 of the homologous peptide antigens (Figure 1A) related to TCR-pMHC complex of SARS-CoV-2 as well as SARS-CoV and bat-CoV N proteins prefers the hydrophobic amino acid residues (e.g. Ile, Leu, Met, and Phe), and the second position of these hit peptides is an hydrophobic amino acid residue Pro forming five strong VDW forces with residues Y99, V67, M45, Y7, and F9 and two H-bonds with residues K66 and E63 on MHC molecule (Figure S2A, left). By contrast, the second position of MERS-CoV N protein-derived hit peptide is charged residue Arg forming three strong VDW forces with residues M45, F9, and V67 and two H-bonds with residues K66 and E63 (As same as SARS-CoV-2) on MHC molecule. Surprisingly, position 9 of all homologous peptides antigens (Figure 1A) prefers the hydrophobic amino acid residues (e.g. Leu, Ile, Val, and Met), and the position 9 of these hit peptides is hydrophobic amino acid residue Leu (in the SARS-CoV-2, SARS-CoV, and bat-CoV) and Gly (in the MERS-CoV). Our results showed that Leu attaches to the MHC with three strong VDW forces with residues L81, I124, and W147 and three H-bonds with residues D77, Y84, and T143, while Gly forms three strong VDW forces with residues L81, V95, and W147 and two H-bonds with residues D77 and V95 on MHC molecule (Figure S2A, left). Moreover, positions 4, 6, and 8 of hit peptides in SARS-CoV-2, SARS-CoV, and bat-CoV form one H-bond with residues Q52, Q52 and D32 in chain E of TCR, respectively (Figure S2C, left). While, only position 4 of hit peptides in MERS-CoV forms one H-bond with reside S100 in chain D of TCR (Table S1). Visualization of interactions in the atomic level structure of a TCR-pMHC complex in the hit peptide of SARS-CoV-2 N protein for HLA-A0201 (Figure S3A) within 20 and 8 Å generated on-the-fly using PyMOL.

**Figure 1.**
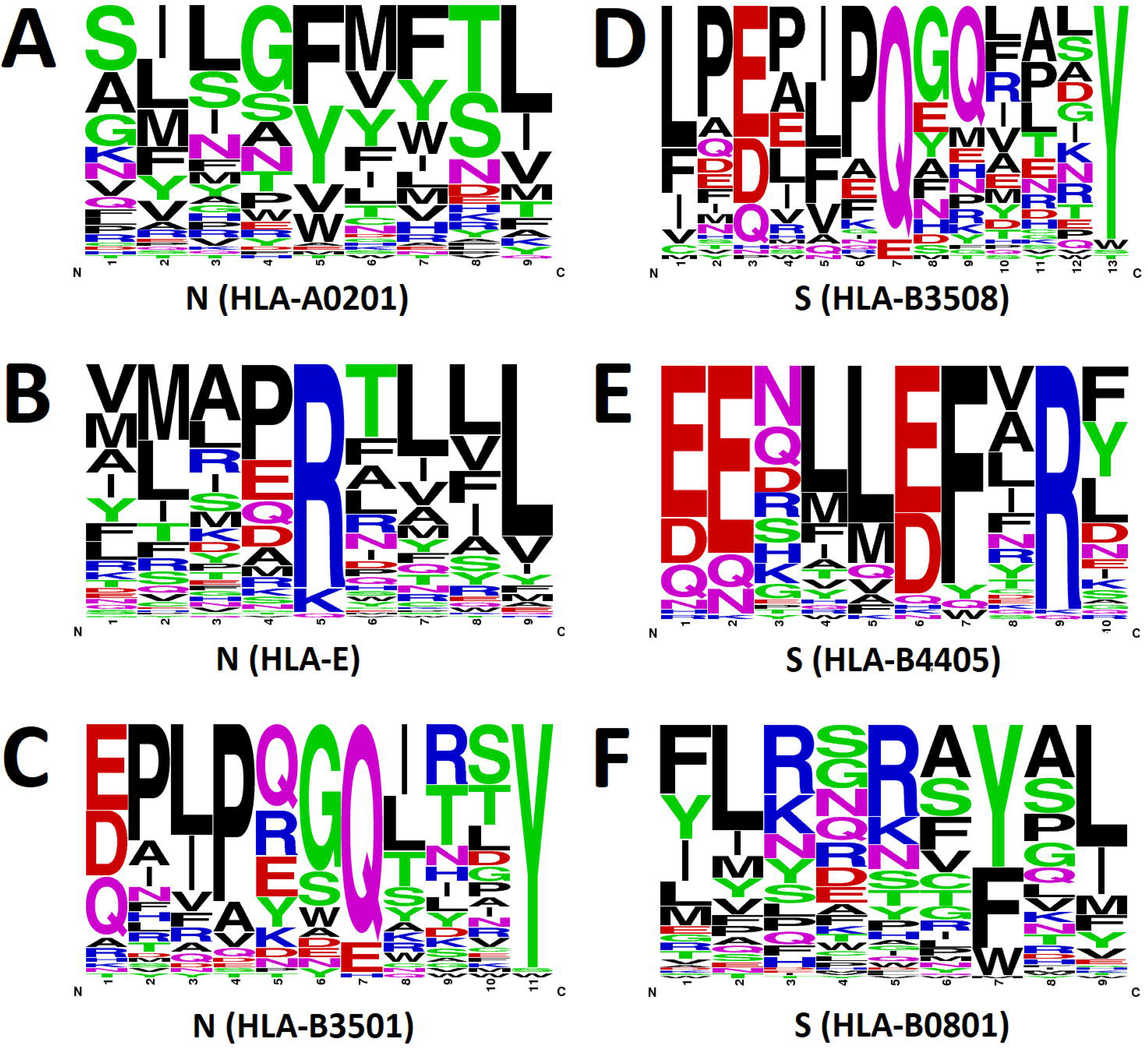
Overview of the amino acid profiles of the homologous peptide antigens related to the SARS-COV-2 N and S proteins associated with HLA-A0201 (A), HLA-E (B), HLA-B3501 (C), HLA-B3508 (D), HLA-B4405 (E), and HLA-B0801 (F) in TCR-pMHC complex template.

### The binding events among hit peptides derived from N proteins related to HLA-E

Our models proposed that positions 1, 2, 7, 8, and 9 of the homologous peptide antigens (Figure 1B) of SARS-CoV-2, bat-CoV, MERS-CoV, and SARS-CoV prefer the hydrophobic amino acid residues. Accordingly, position 1 of these hit peptides forms two H-bonds with residues Y170 and Y158 and position 2 of these hit peptides forms a H-bonds with residue E62 and four strong VDW forces with residues A66, Y6, W96, and M44. Moreover, position 7 of these hit peptides forms a H-bond with residue N76 and three strong VDW forces with residues L123, F115, and W132. (Figure S2A, center). We identified position 4 of hit peptides lacks any contacts, although the position 5 and 6 of hit peptides form both H-bonds and/or strong VDW forces on both MHC molecule and TCR (Figure S2A and S2C, center) (Table S2). Visualization of interactions in the atomic level structure of a TCR-pMHC complex in the hit peptide of SARS-CoV-2 N protein for HLA-E (Figure S3C) within 20 and 8 Å was generated on-the-fly using PyMOL.

### The binding events among hit peptides derived from N proteins related to HLA-B3501

Position 1 of the homologous peptide antigens (Figure 1C) of TCR-pMHC complex of N protein in the SARS-CoV-2, bat-CoV, and SARS-CoV prefers the charged amino acid residues (e.g. Glu and Asp), and the first position of all hit peptides is polar amino acid forming four H-bonds with residues R62, Y171, Y7, and Y159 on MHC molecule (Figure S2A, right) and one H-bond with residue Y96 on TCR (Figure S2C, right). Also, position 2 of the homologous peptide antigens in SARS-CoV-2, SARS-CoV, and bat-CoV prefers the hydrophobic amino acid residues (e.g. Pro, Ala, and Ile) and the position 2 of these hit peptides forms three strong VDW contacts with residues Y159, Y7, and Y99 on MHC molecule (Figure S2A, right). Interestingly, based on the binding models and interactions, positions 6, 8, and 9 of all hit peptides correlated with the SARS-CoV-2, bat-CoV, and SARS-CoV lack any H-bonds and strong VDW contact, while the hit peptides of all comparison targets correlate well with the amino acid profile on the conserved positions (i.e. 4 and 7) forming one strong VDW forces and two H-bonds based on the interactions on TCR (Figure S2C, right) (Table S3). Visualization of interactions in the atomic level structure of a TCR-pMHC complex in the hit peptide of SARS-CoV-2 N protein for HLA-B3501 (Figure S3B) within 20 and 8 Å was generated on-the-fly using PyMOL.

### The binding events among hit peptides derived from S proteins related to HLA-B3508

Position 1 and 11 of the homologous peptide antigens (Figure 1D) in TCR-pMHC complex in all comparison targets, excluding MERS-CoV, prefer the hydrophobic amino acid residues (e.g. Leu, Phe, Ile, Ala, and Pro), and the first position of the hit peptides is residue Leu forming three H-bonds with residues Y159, Y7, and Y171 and two strong VDW contacts with residues W167 and L163 on MHC molecule (Figure S2B, left). Also, position 11 is residue Ala forming one strong VDW contacts with residue W147 on MHC molecule (Figure S2B, left). We found that the position 10 of hit peptides lacks any H-bond and strong VDW forces, while the positions 5 and 6 of hit peptides in the SARS-CoV-2, bat-CoV, and SARS-CoV form one strong VDW forces on MHC molecule (Figure S2B, left) and two strong VDW forces on TCR (Figure S2D, left). Additionally, position 6 forms one H-bond on TCR (Figure S2D, left) (Table S4). Visualization of interactions in the atomic level structure of a TCR-pMHC complex in the hit peptide of SARS-CoV-2 S protein for HLA-B3508 (Figure S3E) within 20 and 8 Å was generated on-the-fly using PyMOL.

### The binding events among hit peptides derived from S proteins related to HLA-B4405

Hit peptide derived from SARS-CoV-2 as same as bat-CoV, MERS-CoV, and SARS-CoV correlates well with the amino acid profile on the conserved positions (i.e. 4, 5, 7, and 8) and all of them form one strong VDW forces with residue I66, V152, and A150 based on the interactions of pMHC interfaces in HLA-B4405-peptide-TCR template (3dxa), respectively. In spite of that, position 9 of all queries forms two H-bonds with residue E76 and W147 on MHC (Figure S2B, center) and one strong VDW forces and two H-bonds on TCR (Figure S2D, center). Based on the interactions of hit peptides with TCR, position 4, 5, and 7 form one, three, and two strong VDW forces, receptivity, Also, position 8 of the homologous peptide antigens (Figure 1E) in all queries prefers the hydrophobic amino acid residues (e.g. Val, Ala, and Leu) and the position 8 of these hit peptides (excluding MERS-CoV) is polar amino acid residue Gln forming one strong VDW contacts with residue V152 on MHC molecule (Figure S2B, center). Whereas position 6 of hit peptides attaches to the TCR molecule with two H-bonds (Figure S2D, center) (Table S5). Visualization of interactions in the atomic level structure of a TCR-pMHC complex in the hit peptide of SARS-CoV-2 S protein for HLA-B4405 (Figure S3F) within 20 and 8 Å was generated on-the-fly using PyMOL.

### The binding events among hit peptides derived from S proteins related to HLA-B0801

As pointed out, positions 2 and 9 of the homologous peptide antigens (Figure 1F) of TCR-pMHC complex related to all queries prefer the hydrophobic amino acid residues (e.g. Leu, Ile, Met) and the second position of these hit peptide in SARS-CoV-2, bat-CoV, and SARS-CoV is polar amino acid residue Ser and in MERS-CoV is hydrophobic amino acid residue Pro forming two strong VDW forces with residues I66 and F36, and one H-bond with residue N63 on MHC molecule (Figure S2B, right). Conversely, position 4 of the homologous peptide antigens of all queries has no detectable binding to both MHC and TCR. Additionally, position 6 of these hit peptides with a H-bond and strong VDW forces connects to the residues Q96 and Y96 on TCR, respectively (Figure S2D, right). The hit peptide correlates well with the amino acid profile on the conserved positions 7 (Tyr) that forms one strong VDW forces with residue W147 on MHC (Figure S2B, right) and four strong VDW forces with residue A97, H47, H32, and L90 on TCR (Figure S2D, right and Table S6). Visualization of interactions in the atomic level structure of a TCR-pMHC complex in the hit peptide of SARS-CoV-2 S protein for HLA-B0801 (Figure S3D) within 20 and 8 Å was generated on-the-fly using PyMOL.

### Immunogenicity study for SARS-CoV-2 protein derived peptide antigens

Prediction the immunogenicity of a class I peptide MHC complex showed that the top putative immunogenic SARS-CoV-2-derived hit peptide candidates related to HLA-A0201 based on their score were 105SPRWYFYYL113 (derived from N protein), 5NVFAFPFTI13 (derived from ORF10 protein), 19LFLAFVVFL27 (derived from E protein), and 145YFLCWHTNC153 (derived from ORF3a protein), respectively. Interestingly, the hit peptides derived from N and E proteins of SARS-CoV-2 by using HLA-A0201 were similar to the peptides derived from N and E proteins of bat-CoV and SARS-CoV, and all of them had higher immunogenicity in comparison to hit peptides derived from MERS-CoV N and E proteins. This result generally was consistent with the result of Grifoni et al., who showed that the sequence of protein consensus of SARS-CoV-2 was similar to the protein sequence of SARS-CoV and bat-CoV, but was more different from protein consensus of MERS-CoV [17]. More specifically, our results showed that the hit peptide derived from ORF3a of SARS-CoV-2 was similar to the hit peptide derived from bat-CoV protein 3, and both of them had higher immunogenicity in comparison to hit peptide derived from MERS-CoV and SARS-CoV ORF3 proteins. Our data emphasized the hit peptide derived from SAR-CoV-2 S protein by using HLA-A0201 had lower immunogenicity than bat-CoV, MERS-CoV, and SARS-CoV. Additionally, regarding HLA-E, comparison of SARS-CoV-2 N-derived hit peptide (DLSPRWYFY) to sequences for bat-CoV (ELSPRWYFY), MERS-CoV (QLAPRWYFY), and SARS-CoV (ELSPRWYFY) revealed a high degree of immunogenicity between SARS-CoV-2, SARS-CoV, and bat-CoV but a more limited immunogenicity with MERS-CoV.

Based on the hypothesis that solvent exposed residues via increasing TCR binding can provide appropriate evidence about peptide immunogenicity, we measured solvent exposed area (SEA) for each hit peptide. Our results displayed that the SEA > 30 Å^2^ for hit peptide related to HLA-B0801, HLA-B3501, and HLA-B3508 were 5.08, 7.76, and 8.28, respectively. Indeed, we found the most solvent accessibility of amino acids were in the hit peptide M (SEA > 30 Å^2^: 5.99), S (SEA > 30 Å^2^: 6.75), and ORF10 (SEA > 30 Å^2^: 6) for HLA-A0201, HLA-B4405, and HLA-E, respectively.

Over the past few months, studies in humans are beginning to unravel the underpinnings relationship between hydrophobicity scales and eradication of immune responses. Currently, identification of peptide regions exposed at the surface has gained much attention in the field of immunogenicity of peptides presented by MHC. This concept is reinforced when we investigate the hydrophobic solvent accessible surface area (hSASA) for each peptide. As such, to account for immunogenicity of MHC-presented peptides, we relied on the hydrophobicity of each peptide position. Further, we evaluated whole-residue hydrophobicity scales among all hit peptides by using the Wimley-White whole-residue hydrophobicity scales to examine how structural characteristics could result in enhanced immunogenicity predictions. Our results are supported by those obtained from IEBD, related to HLA-A0201-peptide-TCR templates showed that specific locations with the most pronounced variations at positions 4, 6, 7, 8, and 9 across the peptides derived from N protein were more hydrophobic in the immunogenic dataset than the peptide derived from M protein (Figure 2A). Our result also showed that the hit peptide antigens derived from N protein related to HLA-E-peptide-TCR (Figure 2E) and S protein related to HLA-B4405 peptide-TCR (Figure 2D) had grater hydrophobicity than ORF10 and M proteins due to differences at positions 6 and 9, and 5, respectively. These observations indicate that hit peptide antigens derived from N and S proteins properly contain immunogenic amino acids. Similarly, positions 8, 10, and 11 across the peptides derived from N, positions 1, 4, 6, 7, and 9 across the peptides derived from S, and positions 1, 4, 5, and 12 across the peptides derived from S proteins (by HLA-B3501 (Figure 2B), HLA-B0801 (Figure 2C), and HLA-B3508 (Figure 2F) had high score of hydrophobicity based on their amino acid sequence, respectively.

**Figure 2.**
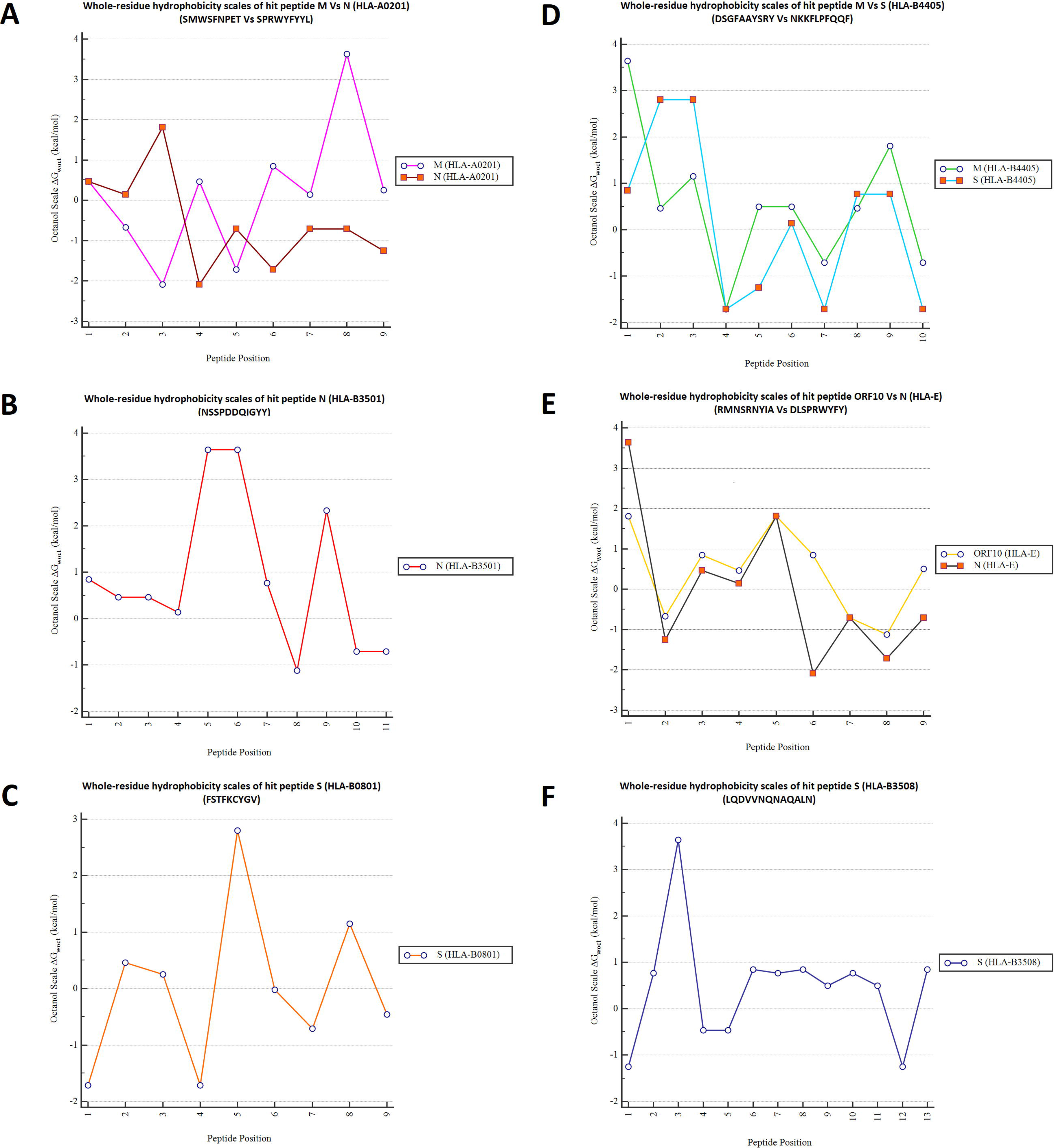
Comparison of the hydrophobicity of each hit peptide position related to SARS-CoV-2 by using the Wimley-White whole-residue hydrophobicity scales. Values less than zero represent greater hydrophobicity in each position of hit peptide. A: Comparison of the hydrophobicity of hit peptide derived from M protein Vs N protein in HLA-A0201; B: Hydrophobicity of hit peptide derived from N protein in HLA-B3501; C: Hydrophobicity of hit peptide derived from S protein in HLA-B0801; D: Comparison of the hydrophobicity of hit peptide derived from M protein Vs S protein in HLA-B4405; E: Comparison of the hydrophobicity of hit peptide derived from ORF10 protein Vs N protein in HLA-E; F: Hydrophobicity of hit peptide derived from S protein in HLA-B03508.

In line with our findings, a landmark study by Grifoni et al. has supporting evidence for the detrimental association between the incidence of CD4^+^ and CD8^+^ response against the N and S proteins in humans with COVID-19 Disease [3]. Consistent with this, there are evidence suggesting that the S protein of SARS-CoV-2 is a promising option for induction of a protective immunity [18]. Sorting this relation out could enhance our understanding of the role of SARS-CoV-2 proteins in the context of T cell immunity. Of note, SARS-CoV-2-derived hit peptide from N and S proteins were selected for further analysis.

### Evaluation of SARS-CoV-2-derived hit peptides sequence conservation

Next, we used CLUSTALW and CLC Sequence Viewer (Version 6.7.1) to calculate the percentage identity between SARS-CoV-2-derived hit peptide sequences and bat-CoV, MERS-CoV, and SARS-CoV. We found that HLA-A0201 binding hit peptide of SARS-CoV-2 N protein (Figure 3A) and the SARS-CoV-2 S proteins correlated with HLA-B3508 (Figure 3F) and HLA-B0801 (Figure 3B) were more strongly conserved between bat-CoV and SARS-CoV (100% identity) in comparison with MERS-CoV (77.78%, 69.23%, and 22.22% identity, respectively). We also identified that the hit peptide of SARS-CoV-2 N protein correlated to HLA-B3501 (Figure 3E) and HLA-E (Figure 3C) had 45.45% and 77.78% identity with MERS-CoV, and 90.91% and 88.89% with bat-CoV and SARS-CoV, respectively. Correspondingly, HLA-B4405 binding hit peptide of SARS-CoV-2 S protein (Figure 3D) had 25% identity with MERS-CoV, 60% identity with bat-CoV, and 70% identity with SARS-CoV, suggesting high sequences similarity between hit peptides derived from N and S proteins of SARS-CoV-2, SARS-CoV, and bat-CoV compared to MERS-CoV.

**Figure 3.**
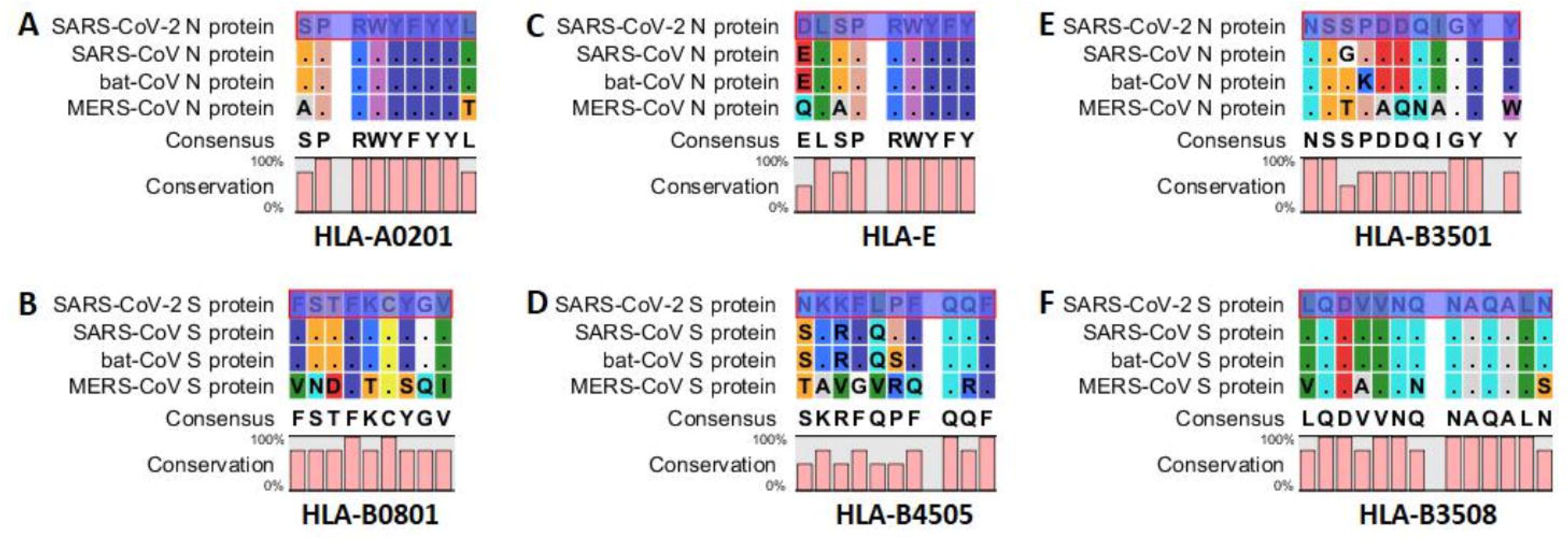
Amino acid sequence alignment of hit peptides derived from N and S proteins related to SARS-CoV-2, SARS-CoV, bat-CoV, and MERS-CoV by using the program CLC Sequence Viewer (Version 6.7.1). Upper panel: Amino acid sequence alignment of hit peptides derived from N proteins in HLA-A0201 (A), HLA-E (C), and HLA-B3501 (E); Bottom panel: Amino acid sequence alignment of hit peptides derived from S proteins in HLA-B0801 (B), HLA-B4405 (D), and HLA-B3508 (F).

### Assessment of residues interactions

It has long been known that a potent immune response to hit peptides is mediated by the higher affinity and binding of the hit peptides to immune cells under optimal conformation. Better understanding of specific macromolecular interactions within the binding sites could help identifying molecular recognitions such as those occurringamong targets and ligands, which are important for drug development. This knowledge is expected to open new application avenues for the designing ofeffective vaccines based on high prioritized epitopes that have strong affinity and binding to TCR and MHC. Different SARS-CoV-2-derived hit peptides with immunogenicity prediction displayed that the maximum binding rate of chain C (hit peptide) in TCR-pMHC complex was linked to the N and S proteins, respectively. Apart from MERS-CoV, when we compared the hydrophobic contacts, mutually exclusive interactions, and polar contacts among different hit peptides by using HLA-A0201- (Figure S4A), HLA-B0801- (Figure S4B), HLA-B3508- (Figure S4D), and HLA-E-peptide-TCR template (Figure S4F), it was proved that these interactions were equal in chain C of TCR-pMHC complex associated with N and S proteins of SARS-CoV-2, bat-CoV, and SARS-CoV proteins. The above-mentioned interactions were lower in SARS-CoV-2 than bat-CoV and SARS-CoV associated with HLA-B3501 (Figure S4C) similarly to the hydrophobic contacts by using HLA-E (Figure S4F). By contrast, the number of aforementioned contents (excluding mutually exclusive interactions in SARS-CoV by using HLA-B4405 (Figure S4E)) was higher in SARS-CoV-2 than bat-CoV and SARS-CoV associated with HLA-B3508 (Figure S4D) as well as the number of polar contacts by HLA-E (Figure S4F). Correspondingly, in HLA-E, SARS-CoV-2-derived hit peptide had the lower mutually exclusive interactions compared to the bat-CoV and SARS-CoV. On the opposite side, this type of interatomic forces was higher in the SARS-CoV-2-derived hit peptide than MERS-CoV associated with HLA-E (Figure S4F, Table 2, and Table S7-S9).

**Table 2:** Summary of the interatomic interactions within chain C and their binding site interactions in the TCR-pMHC complex of SARS-CoV-2 proteins

| Interatomic interactions |  | HLA-A0201 |  |  |  |  | HLA-B0801 | HLA-B3501 | HLA-B3508 | HLA-B4405 |  | HLA-E |  |  |  |
| --- | --- | --- | --- | --- | --- | --- | --- | --- | --- | --- | --- | --- | --- | --- | --- |
|  |  | M | E | N | ORF3a | ORF10 | S | S | N | S | M | S | N | ORF10 | S |
| Mutually Exclusive Interactions | VDW interactions | 7 | 12 | 17 | 13 | 12 | 5 | 9 | 12 | 16 | 8 | 16 | 32 | 27 | 32 |
|  | VDW clash interactions | 27 | 46 | 100 | 60 | 36 | 36 | 64 | 39 | 56 | 29 | 49 | 130 | 58 | 92 |
|  | Covalent interactions | 0 | 1 | 7 | 2 | 0 | 0 | 3 | 0 | 1 | 0 | 1 | 6 | 1 | 5 |
|  | Covalent clash interactions | 0 | 0 | 1 | 0 | 0 | 0 | 1 | 0 | 0 | 0 | 0 | 2 | 0 | 0 |
|  | Proximal | 675 | 738 | 878 | 777 | 708 | 646 | 640 | 669 | 926 | 667 | 791 | 883 | 748 | 816 |
|  | Total | 709 | 797 | 1003 | 852 | 756 | 687 | 717 | 720 | 999 | 704 | 857 | 1053 | 834 | 945 |
| Polar Contacts | Polar contacts | 18 | 12 | 16 | 23 | 14 | 16 | 20 | 15 | 23 | 15 | 17 | 24 | 24 | 20 |
|  | Weak polar contacts | 10 | 18 | 31 | 16 | 15 | 17 | 23 | 13 | 26 | 17 | 22 | 43 | 29 | 39 |
|  | Total | 28 | 30 | 47 | 39 | 29 | 33 | 43 | 28 | 49 | 32 | 39 | 67 | 53 | 59 |
| Feature Contacts | Hydrogen bonds | 16 | 13 | 15 | 16 | 14 | 13 | 16 | 10 | 18 | 11 | 14 | 16 | 16 | 16 |
|  | Weak hydrogen bonds | 11 | 12 | 13 | 11 | 15 | 13 | 9 | 11 | 10 | 8 | 6 | 14 | 13 | 13 |
|  | Ionic interactions | 0 | 0 | 0 | 1 | 0 | 0 | 0 | 2 | 0 | 5 | 2 | 1 | 0 | 0 |
|  | Aromatic contacts | 2 | 10 | 40 | 14 | 6 | 0 | 4 | 2 | 0 | 7 | 7 | 99 | 0 | 49 |
|  | Hydrophobic contacts | 49 | 125 | 99 | 76 | 82 | 66 | 86 | 34 | 62 | 41 | 90 | 142 | 46 | 96 |
|  | Carbonyl interactions | 1 | 1 | 1 | 1 | 1 | 1 | 0 | 1 | 2 | 0 | 0 | 0 | 0 | 0 |
|  | Total | 79 | 161 | 168 | 119 | 118 | 93 | 115 | 60 | 92 | 72 | 119 | 272 | 75 | 203 |

Chain C: Hit peptide; VDW: van der Waals (VDW) forces; E: Envelope protein; M: Membrane glycoprotein; N: Nucleocapsid phosphoprotein; ORF: Open reading frame; S: Surface glycoprotein; HLA: Human leukocyte antigen

### Peptide-protein flexible Docking

For various kinds of biological applications such as vaccine design, peptides have been introduced as potential candidates. Recently, several methods have been introduced for protein-peptide docking, which can estimate the structure of the protein-peptide complex through peptide sequence and protein structure. Accordingly, for hit peptides derived from N protein, NSSPDDQIGYY had the highest interaction similarity score with HLA-B3501 than SPRWYFYYL and DLSPRWYFY with HLA-A0201 and HLA-E, respectively. Moreover, for hit peptides derived from S protein, FSTFKCYGV had the highest interaction similarity score with HLA-B0801 than LQDVVNQNAQALN and NKKFLPFQQF with HLA-B3508 and HLA-B4405, respectively. Figure 4 represents a sample of successful docking model between each hit peptide and its MHC protein.

**Figure 4.**
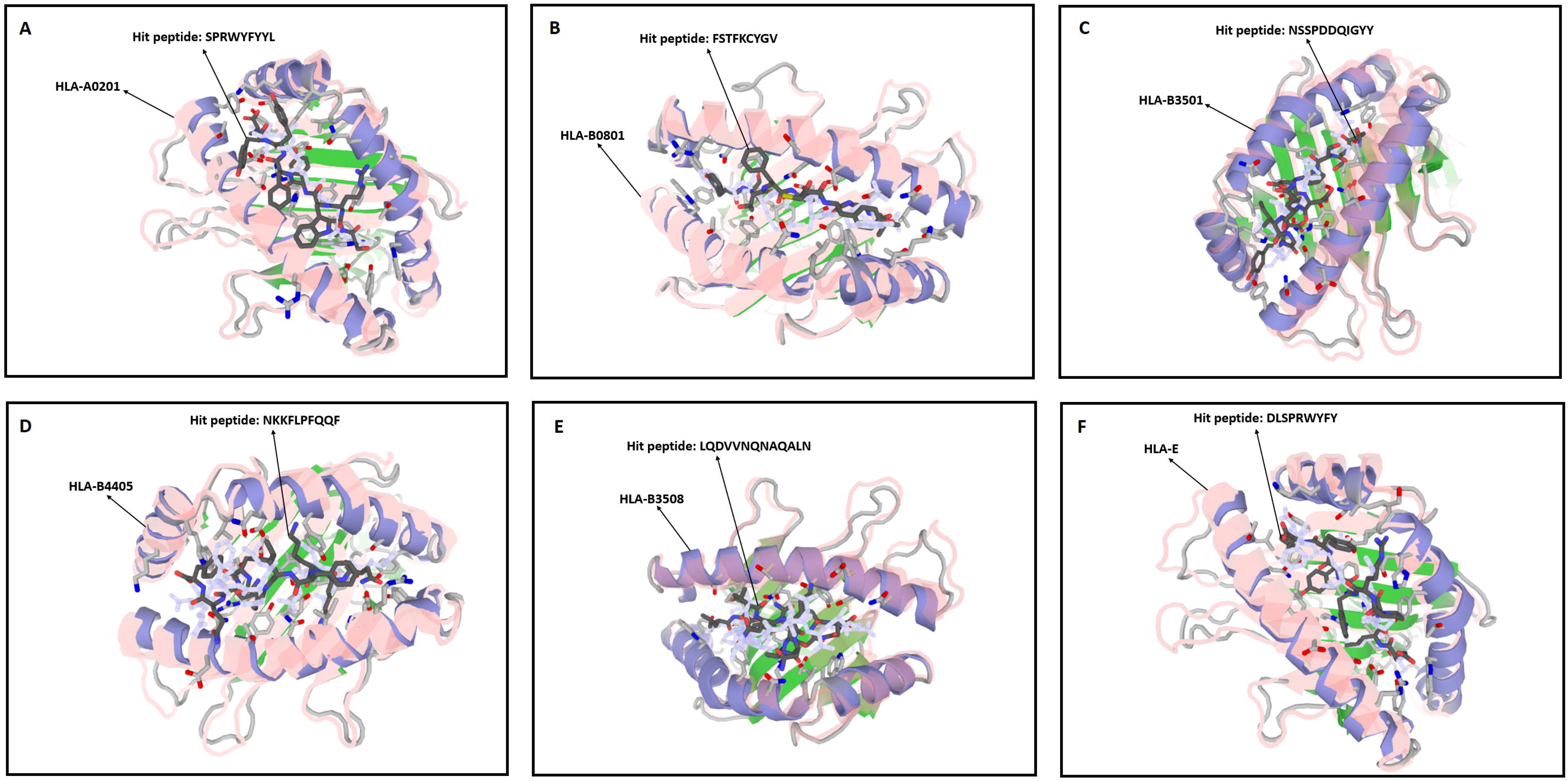
A: Successful peptide-protein docking between hit peptide derived from N protein (SPRWYFYYL) and HLA-A0201 with cluster density scores of 219; B: Successful peptide-protein docking between hit peptide derived from S protein (FSTFKCYGV) and HLA-B0801 with cluster density scores of 272; C: Successful peptide-protein docking between hit peptide derived from N protein (NSSPDDQIGYY) and HLA-B3501 with cluster density scores of 270; D: Successful peptide-protein docking between hit peptide derived from S protein (NKKFLPFQQF) and HLA-B4405 with cluster density scores of 178; E: Successful peptide-protein docking between hit peptide derived from S protein (LQDVVNQNAQALN) and HLA-B3508 with cluster density scores of 209; F: Successful peptide-protein docking between hit peptide derived from N protein (DLSPRWYFY) and HLA-E with cluster density scores of 215.

### Multi-hit peptides vaccine and its structural properties

On the basis of immunogenicity and hydrophobicity scale, and residues interactions, six hit peptides derived from N and S proteins related to HLA-A0201, HLA-B0801, HLA-B3501, HLA-B3508, HLA-B4405, and HLA-E including SPRWYFYYL, FSTFKCYGV, NSSPDDQIGYY, LQDVVNQNAQALN, NKKFLPFQQF, and DLSPRWYFY were joined together by linking the N and S hit peptides via AAY and GPGPG linkers. Additionally, the human beta defensin 3 was also attached to the N-terminus of the finalized vaccine construct using EAAAK linker which acts as adjuvant to improve the immunogenicity of the vaccine. Evaluation of the final vaccine construct amino acid sequences (comprising a length of 122 amino acids) by BLASTp showed only 36% similarity which is due to the human β-defensin 3 sequence, proving this vaccine did not have any overlap with human proteins and could not elicit autoimmunity. Assessment the secondary structural of the vaccine using SOPMA server uncovered that it was composed of 17.21 % Alpha helix (Hh), 24.59 % Extended strand (Ee), 3.28 % Beta turn (Tt), and 54.92 % Random coil (Cc). The allergenicity and antigenicity of the finalized vaccine construct showed that the vaccine is non-allergenic and it has the antigenicity likelihood of 0.4838 with threshold of 0.4, characterizing the antigenic nature of our final multi-hit peptides vaccine for an adequate inducing of immune response. The molecular weight of vaccine construct and Theoretical protrusion index (pI) score was found to be 13498.44 Da and 9.51 while the total numbers of negative and positive charge residues were 7 and 18 respectively, confirming vaccine tendency to be antigenicity and its slightly acidic nature. Indeed, the estimated half-life was 30 hours (mammalian reticulocytes, in vitro), > 20 hours (yeast, in vivo), and >10 hours (*Escherichia coli*, *in vivo*). The extinction coefficient was estimated to be 19285 M^−1^ cm^−1^, at 280 nm measured in water, assuming all pairs of cysteine residues form cystines. The instability index (II) was computed to be 47.60, showing unstable nature of vaccine in the experimental setup. Aliphatic index was calculated as 56.8 while Grand average of hydropathicity (GRAVY) was calculated −0.593.

### Tertiary structure, refinement, and epitope mapping

The three-dimensional (3D) structures supply vital insights into the tertiary structure and position of the hit peptides [19]. In the first step, the 3D structure of the multi-hit peptides vaccine with total 122 amino acid residues were modeled by using the I-TASSER web tool (Figure 5A) Accordingly, 5 final models were predicted as a single domain without disorder. Among them, the first model had a better quality in most cases with C-score = −3.25, Estimated TM-score = 0.35±0.12, and Estimated RMSD = 11.7±4.5Å. However, because of the modelling errors and unavailability of an appropriate template such as angles and irregular bonds, generation of the 3D models cannot be sufficient to follow the necessary accuracy level for some biological purpose, especially where experimental data is rare. As such, for modification of local errors, helping to bring 3D model of vaccine closer to native structures, and growing the accuracy of primary 3D model, the refinement of 3D structure of the vaccine is vital, particularly for furthering in-silico studies [20]. Therefore, the refinement of the model was performed by using 3D^refine^ tool. On the basis of the overall quality of the refined model, the model 5 exhibited the best results with RMSD 0.549Å (Figure 5B). The quality of the best model of the multi-hit peptides vaccine construct was validated by ProSA-web. It is well-known that the Z score is in relation with the length of the protein, indicating that negative Z-scores are more appropriate for a trustworthy model. In fact, the Z score shows the overall quality and calculates the deviation of the total energy related to the protein structure [21]. The Z score of the protein under study was displayed in the plot with a dark black point. Accordingly, our results revealed the improvement of the refined model of vaccine (Z score: −3.81) (Figure 5D) in comparison with initial model (Z score: 3.45) (Figure 5C), which was lying inside the scores range of the native proteins of similar size (range −20 to 10) and also was located within the space of protein related to X-ray, suggesting that the obtained model is reliable and closes to experimentally determined structure. Evaluation the overall quality of the finalized model of vaccine construct by Ramachandran plot analysis emphasized that 77.5% (93/120) of all residues in finalized model (Figure 5F) compared to 54.2% (65/120) of all residues in initial model (Figure 5E) were in favored (98%) regions, also, 90.0% (108/120) of all residues in finalized model (Figure 5F) compared to 78.3% (94/120) of all residues in initial model (Figure 5E) were in allowed (>99.8%) regions. After that, protein structure and visualization of the measured interactions between atoms including the strongest mutually exclusive interactions, polar contacts, H-bonds, ionic interactions, aromatic contacts, hydrophobic contacts, and carbonyl interactions were compared together and analyzed by Arpeggio server and finally the results were tabulated in Table 3.

**Table 3:** Summary of the interatomic interactions in the vaccine constract

| Interatomic interactions |  | Vaccine<br>(Before refinement) | Vaccine<br>(After refinement) |
| --- | --- | --- | --- |
| Mutually<br>Exclusive<br>Interactions | VDW interactions | 166 | 150 |
|  | VDW clash interactions | 419 | 326 |
|  | Covalent interactions | 2 | 2 |
|  | Covalent clash interactions | 0 | 0 |
|  | Proximal | 5479 | 5086 |
|  | Total | 6060 | 5564 |
| Polar<br>Contacts | Polar contacts | 187 | 207 |
|  | Weak polar contacts | 210 | 178 |
|  | Total | 397 | 385 |
| Feature<br>Contacts | Hydrogen bonds | 140 | 137 |
|  | Weak hydrogen bonds | 77 | 76 |
|  | Ionic interactions | 28 | 23 |
|  | Aromatic contacts | 40 | 34 |
|  | Hydrophobic contacts | 327 | 316 |
|  | Carbonyl interactions | 41 | 35 |
|  | Total | 653 | 621 |

VDW: van der Waals (VDW) forces

**Figure 5:**
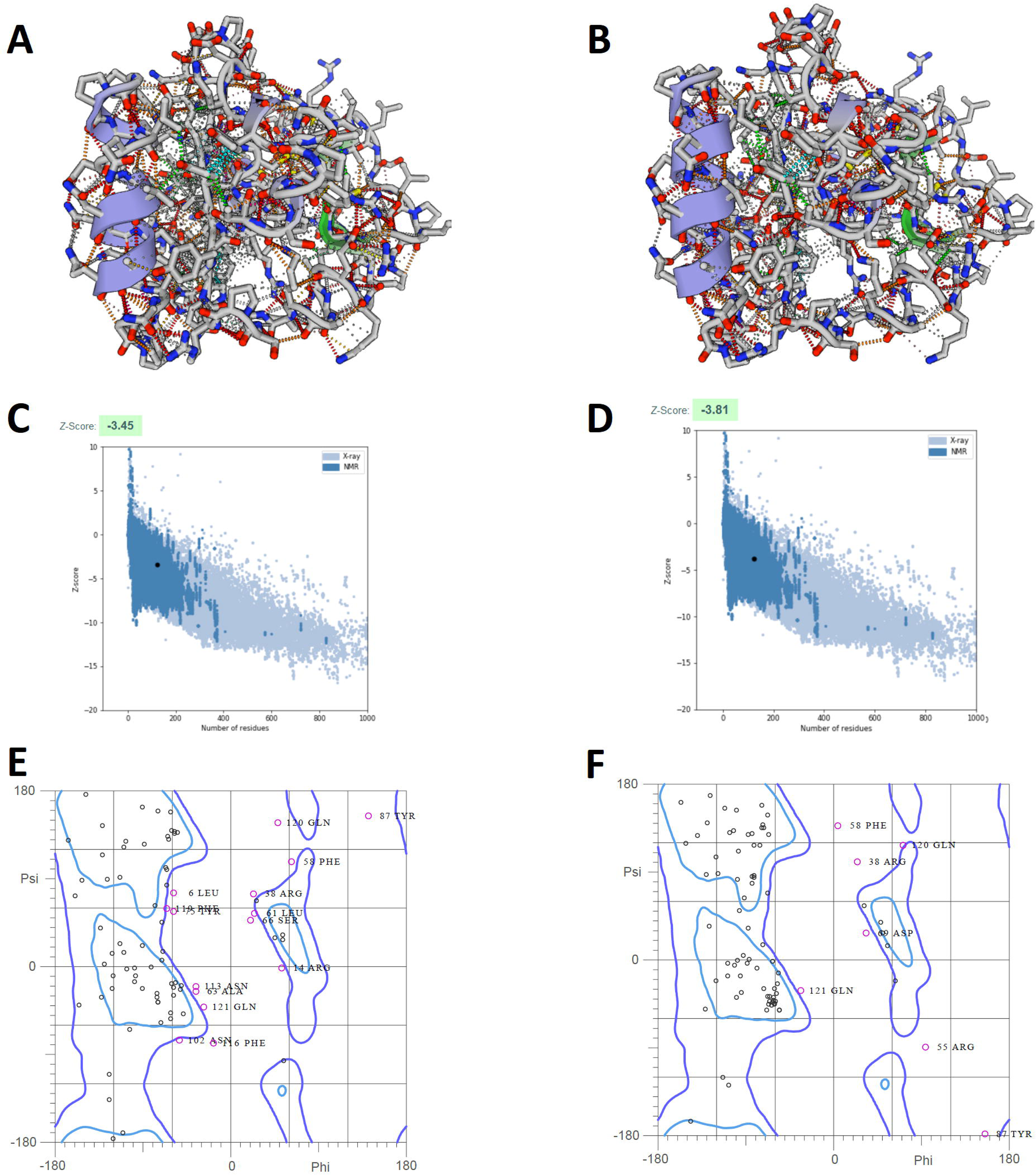
A and B: Schematic structure of the predicted model of the vaccine: The 3D structure of the designed vaccine was suggested through homology modeling by I-TASSER with C-score = −3.25, Estimated TM-score = 0.35±0.12, and Estimated RMSD = 11.7±4.5Å (C-score shows the confidence of model, Estimated TM-score and RMSD are based on C-score and protein length following the correlation discovered between these qualities), then the best proposed model was refined by 3D^refine^. On the basis of the overall quality of the refined model, the model 5 exhibited the best results with RMSD 0.549 Å. Higher score indicates aggressive refinement. The figure 5A and B are in symmetry with the information provided in Table 3 and showing the interacting residues; C and D: ProSA-web Z-score plot. ProSA-web Z-score observed by NMR spectroscopy (dark blue) and X-ray crystallography (light blue) based on the length. The Z-score of the model before (C) and after (D) refinement were −3.45 and −3.81, respectively, which is in the range of native protein conformations; E and F: Ramachandran plot of the modeled vaccine construct before (E) and after (F) of refinement: Ramachandran plot of the model before refinement showed that 54.2% and 78.3% of residues were located in the favored, allowed, respectively (E). While, Ramachandran plot of the model after refinement showed that 77.5% and 90% of residues were located in the favored, allowed, respectively (F).

### Immune simulation

C-ImmSim Immune simulator web server was used for determining ability of vaccine to induce T cell immunity. This server yielded results consistent with actual immune responses as evidenced by a general marked increase in the generation of secondary responses. For better following the effects of the final vaccine construct for stimulation of T cell immunity, a construct with point mutations on key residues, replacing of the hydrophobic amino acids with charged amino acids was constructed (Figure 6A). According to data, after vaccination with native vaccine, there was a consistent rise in Th (helper) cell population with memory development compared to control (Figure 6B-E). Interestingly, immune simulation results demonstrated that while a notable activation of TC (cytotoxic) was observed after vaccination with native vaccine, an activation was not observed after vaccination with the mutant vaccine, proving effectiveness of the native vaccine construct for induction of T cell immune response. (Figure 6F and G). The results definitely showed that the T cell population was highly responsive as the memory response developed.

**Figure 6:**
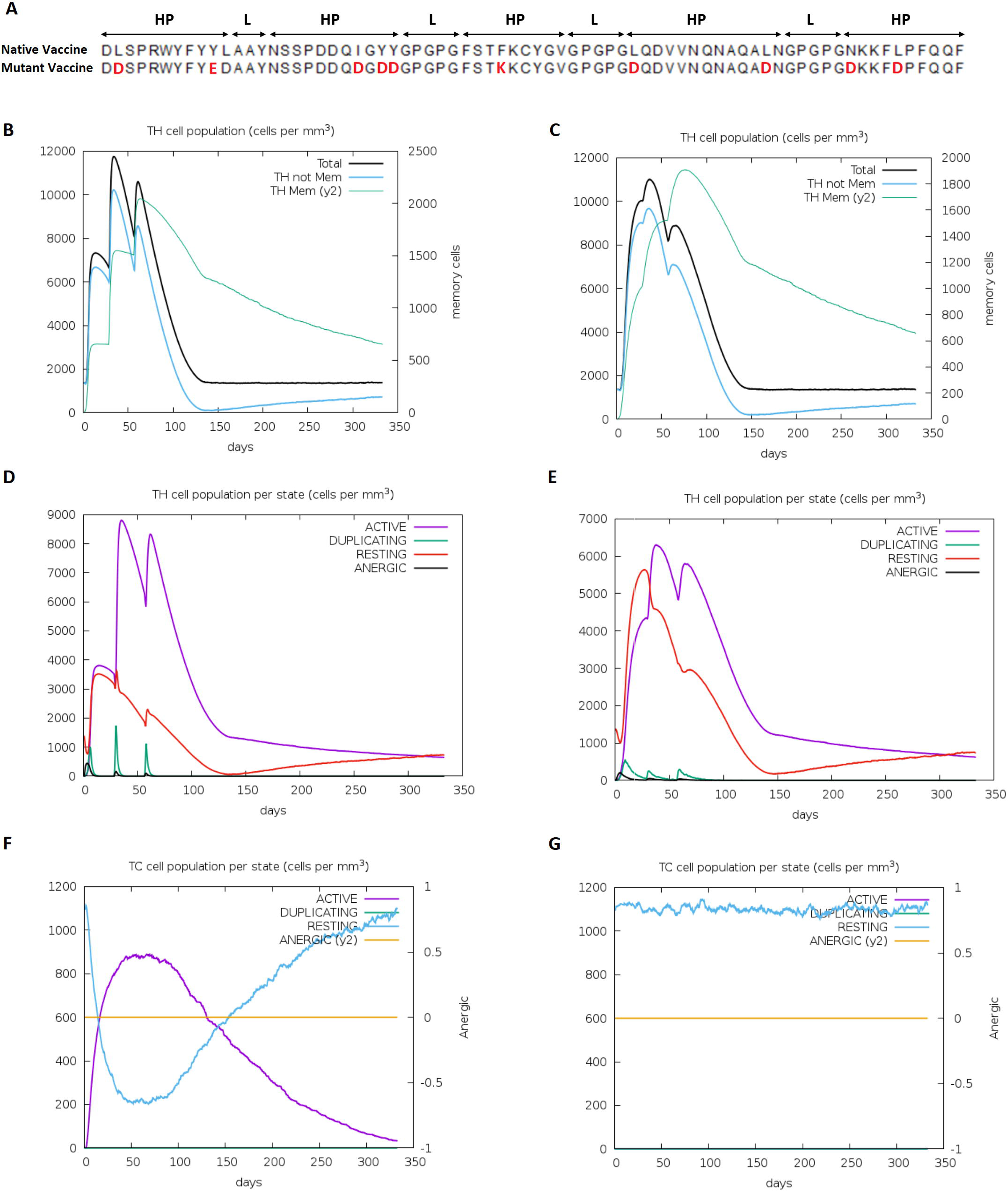
In-silico T cell immune response simulation after administering native and mutant vaccine construct. A: Alignment between native and mutant vaccine construct (HP: hit peptide and L: linker). B-E: CD4 T-helper lymphocytes count after vaccination with native (B and D) and mutant construct (C and E): The plot displays total and memory counts. F-G: CD8 T-cytotoxic lymphocytes count after vaccination with native (F) and mutant construct (G). The plot shows total counts.

## Conclusion

To our best knowledge, from the standpoint of immunoinformatics approaches, the concept discussed in this study is the first structural modeling to investigate both the TCR and pMHC interfaces for the SARS-CoV-2 proteins. Our results provide a blueprint for inferring the SARS-CoV-2-derived hit peptides with high accuracy towards vaccine development. Therefore, it is tempting to speculate that the aforementioned models will offer valuable framework for identifying specific peptides with a potential to acrivate T cell-mediated immune responses to target SARS-CoV-2. As a proof of concept, our study denotes a landmark in the advancement of immunoinformatics based method for designing a multi-hit peptides vaccine against the S and N protein of SARS-CoV-2. In addition, our innovative vaccine candidates can serve as promising tools towards fighting SARS-CoV-2 infections. Further *in vitro* and *in vivo* validation studies are necessary to confirm the safety and efficacy of T-cell-based immunotherapies using our vaccine proposed SARS-CoV-2 peptides.

## Declarations

### Availability of data and material (data transparency)

Not applicable

### Code availability (software application or custom code)

Not applicable

### Authors’ contributions

Amir Hossein Mohseni and Sedigheh Taghinezhad-S contributed equally to this work. Also, all authors discussed the results, commented on the manuscript, and approved the final manuscript.

## Figure Legends

Figure S1. Overall structure of TCR-pMHC complex of SARS-CoV-2 proteins linked to HLA-A0201 (A), HLA-B3501 (B), HLA-B4405 (C), HLA-B0801 (D), HLA-B3508 (E), and HLA-E (F). Hit peptides (Chain C), TCR (Chain E), TCR (Chain D), and MHC (Chain A) are colored purple, blue, red, and green, respectively.

Figure S2: Schematic representation of binding events of both p-MHC and p-TCR interfaces. Atomic binding models with H-bonds are represented by pink and blue lines, respectively. Also, VDW forces are presented. TCR chain D: yellow spiral lines, TCR chain E: pink spiral lines, MHC chain A: blue spiral lines.

Figure S3. Graphic illustration of the interatomic contacts of SARS-CoV-2-derived hit peptide within TCR-pMHC complex for N and S proteins associated with various HLA by using PyMOL. MHC (chain A), hit peptide (chain C), and TCR (chain D and E) are represented by blue, red, yellow, and green color, respectively. Dashed bonds are used to visualize interatomic interactions of hit peptide within TCR-pMHC complex. VDW interactions and VDW clash interactions: gray, H-bonds: red, Weak H-bonds: orange, Aromatic contacts: blue, Hydrophobic contacts: green, Carbonyl interactions: light magenta. Distance of the interaction, overlapping the radii of VDW, and interactions associated with proximal is denoted by the thickness, thickest, and thinnest dashes, respectively.

Figure S4. WebGL-based visualization of interatomic interactions between chain C (hit peptide) and their TCR and MHC molecules belong to the SARS-CoV-2 compared to SARS-CoV, Bat-CoV, and MERS-CoV for HLA-A0201 (A), HLA-B0801 (B), HLA-B3501(C), HLA-B3508 (D), HLA-B4405 (E), and HLA-E (F).

