## Supplementary figures and images for "Inferring MHC interacting SARS-CoV-2 epitopes recognized by TCRs towards designing T cell-based vaccines"

### Figure S1

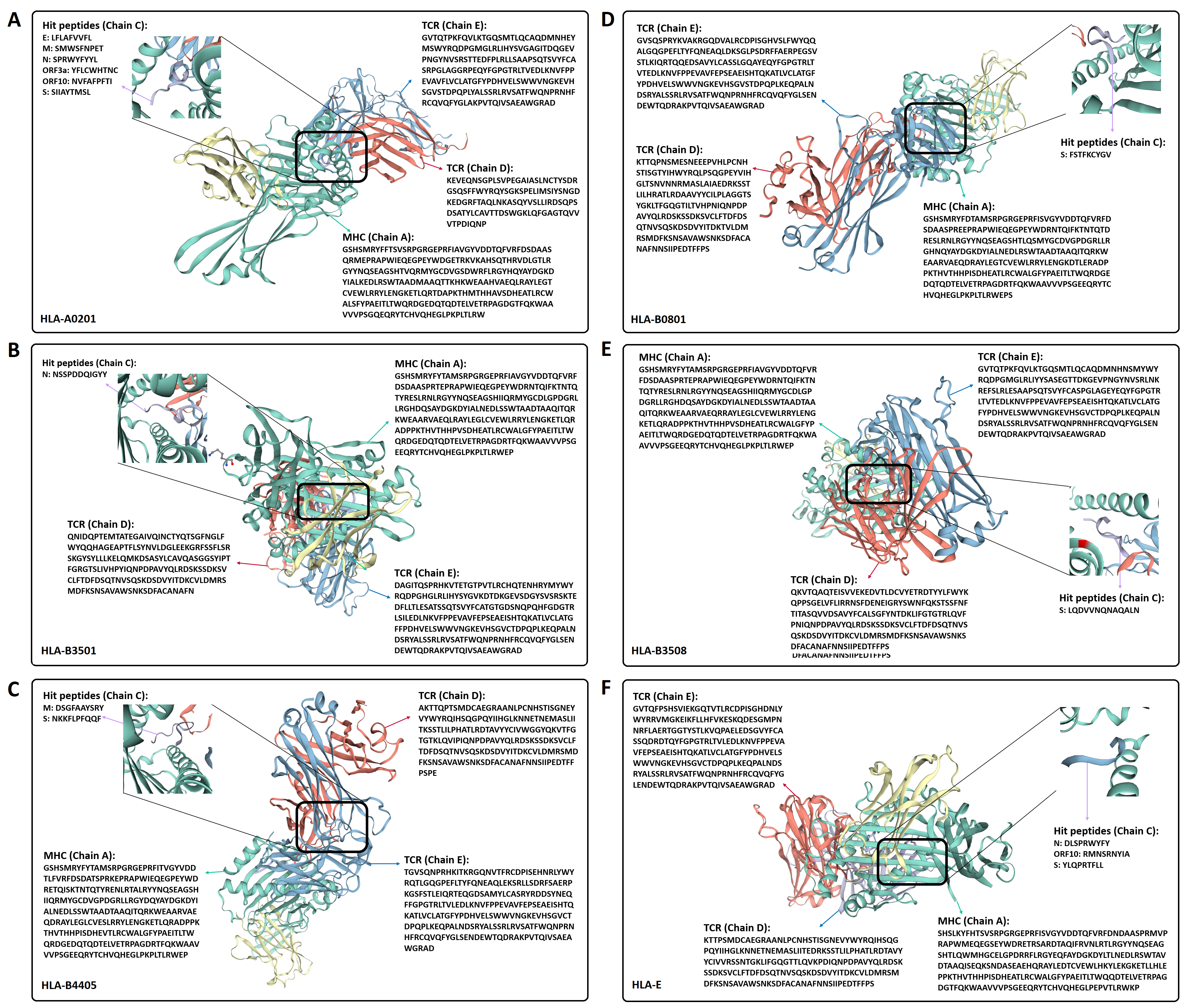

### Figure S2

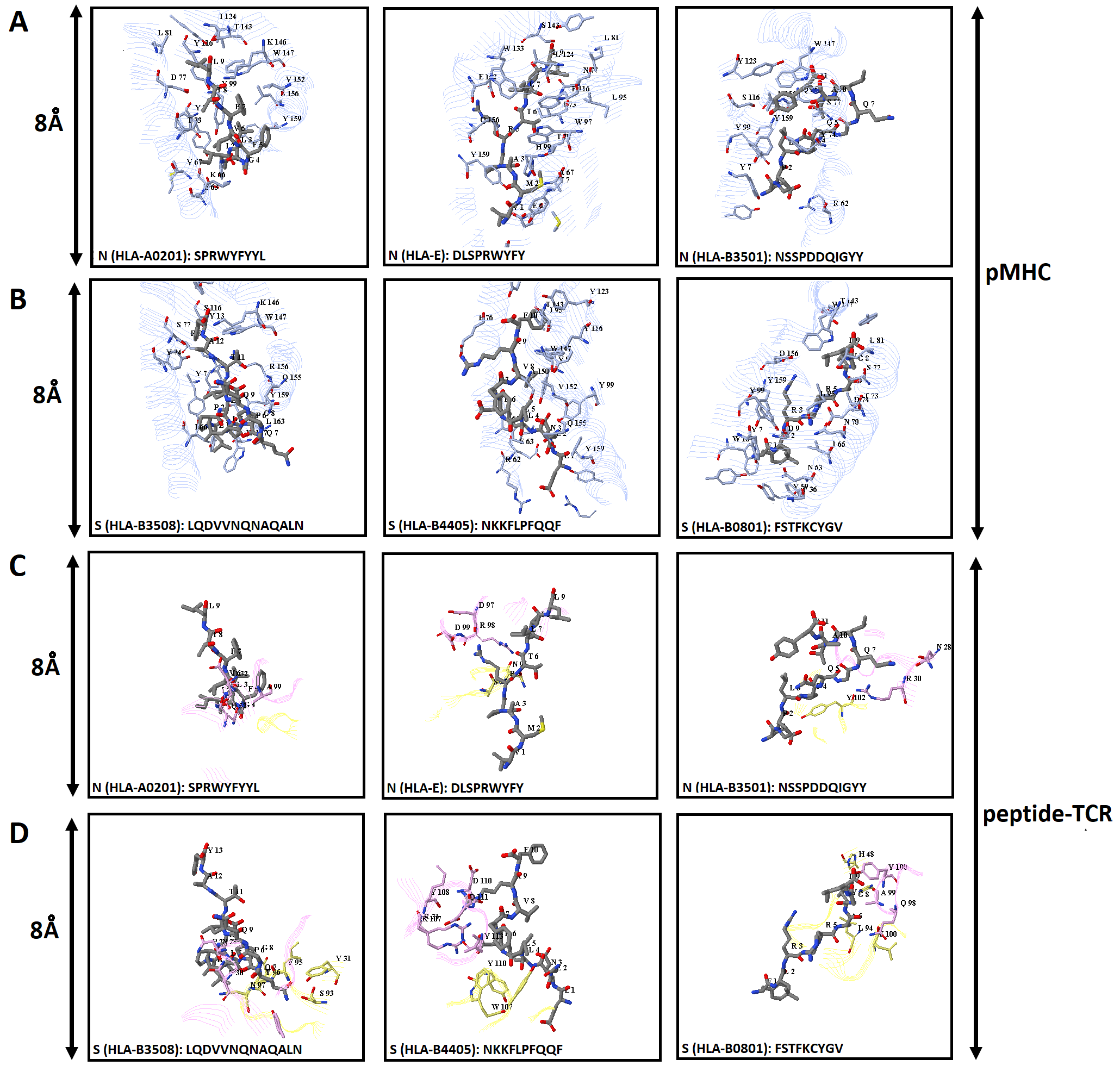

### Figure S3

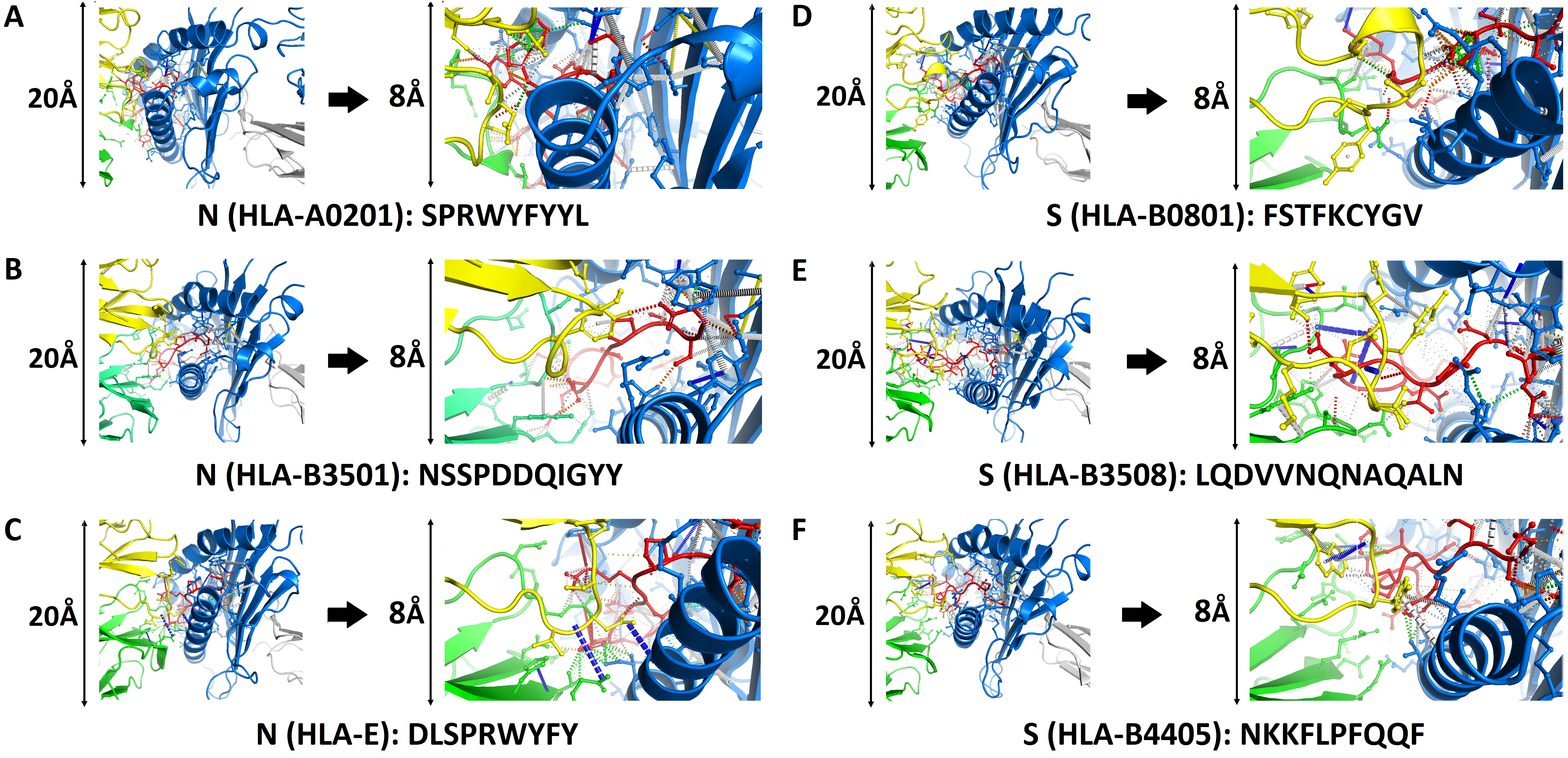

### Figure S4

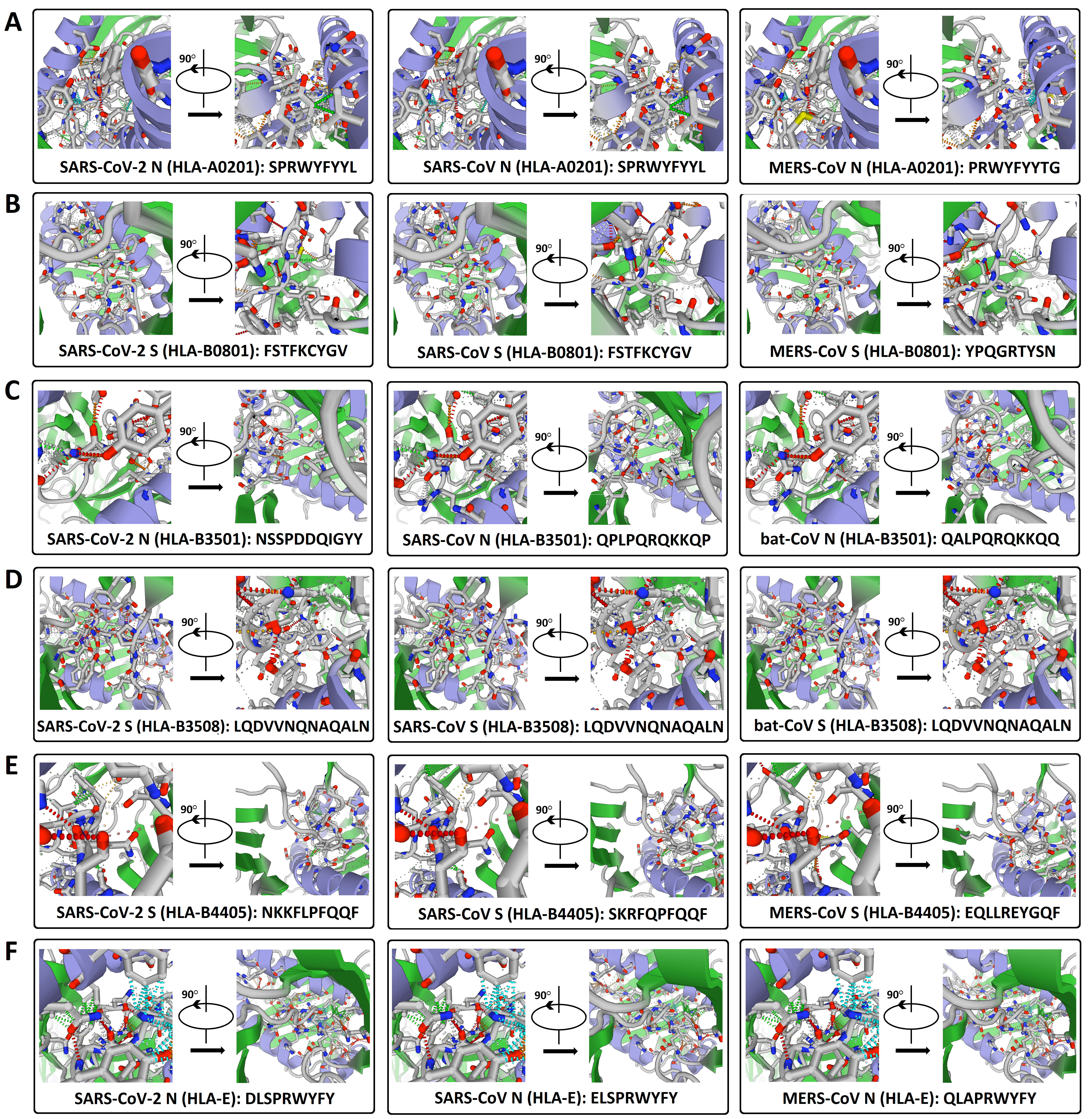
