## Supplementary material for "Inferring MHC interacting SARS-CoV-2 epitopes recognized by TCRs towards designing T cell-based vaccines": Table S1

| **Table S1: Detailed interacting information of the hit peptide antigens of the N query of SARS-CoV-2 in models of TCR-pMHC complex for HLA-A0201** | | | | | |
| --- | --- | --- | --- | --- | --- |
| **Peptide  amino acid** | **MHC molecule** | | **TCR molecules** | | |
|  | **Amino acid** | **Bond type** | **Chain** | **Amino acid** | **Bond type** |
| S | Y159 | H-bond | - | - | - |
|  | Y171 | H-bond |  |  |  |
|  | Y7 | H-bond |  |  |  |
| P | Y99 | Strong VDW | - | - | - |
|  | V67 | Strong VDW |  |  |  |
|  | K66 | H-bond |  |  |  |
|  | M45 | Strong VDW |  |  |  |
|  | Y7 | Strong VDW |  |  |  |
|  | E63 | H-bond |  |  |  |
|  | F9 | Strong VDW |  |  |  |
| R | Y99 | H-bond | - | - | - |
|  | L156 | Strong VDW |  |  |  |
| W | - | - | E | Q52 | H-bond |
| Y | L156 | Strong VDW | - | - | - |
| F | - | - | E | A99 | Strong VDW |
|  |  |  | E | Q52 | H-bond |
| Y | L156 | Strong VDW | - | - | - |
|  | V152 | Strong VDW |  |  |  |
|  | Y116 | Strong VDW |  |  |  |
|  | W147 | Strong VDW |  |  |  |
| Y | T73 | Strong VDW | E | D32 | H-bond |
|  | W147 | H-bond |  |  |  |
|  | K146 | H-bond |  |  |  |
| L | L81 | Strong VDW | - | - | - |
|  | D77 | H-bond |  |  |  |
|  | Y84 | H-bond |  |  |  |
|  | T143 | H-bond |  |  |  |
|  | I124 | Strong VDW |  |  |  |
|  | W147 | Strong VDW |  |  |  |
| VDW: van der Waals (VDW) forces; H- bond: hydrogen bonds | | | | | |
