## Supplementary material for "Inferring MHC interacting SARS-CoV-2 epitopes recognized by TCRs towards designing T cell-based vaccines": Table S2

| **Table S2: Detailed interacting information of the hit peptide antigens of the N query of SARS-CoV-2 in models of TCR-pMHC complex for HLA-E** | | | | | |
| --- | --- | --- | --- | --- | --- |
| **Peptide**  **amino acid** | **MHC molecule** |  | **TCR molecules** |  |  |
|  | **Amino acid** | **Bond type** | **Chain** | **Amino acid** | **Bond type** |
| D | Y170 | H-bond | - | - | - |
|  | Y158 | H-bond |  |  |  |
| L | A66 | Strong VDW | - | - | - |
|  | Y6 | Strong VDW |  |  |  |
|  | W96 | Strong VDW |  |  |  |
|  | M44 | Strong VDW |  |  |  |
|  | E62 | H-bond |  |  |  |
| S | W96 | Strong VDW | - | - | - |
|  | Y158 | Strong VDW |  |  |  |
|  | H98 | Strong VDW |  |  |  |
| P | - | - | - | - | - |
| R | Q155 | H-bond | D | S92 | H-bond |
|  | E151 | H-bond | E | D97 | H-bond |
|  | W96 | Strong VDW | E | D95 | H-bond |
| W | T69 | Strong VDW | D | N93 | H-bond |
|  | W96 | Strong VDW | E | R96 | H-bond |
| Y | L123 | Strong VDW | - | - | - |
|  | F115 | Strong VDW |  |  |  |
|  | N76 | H-bond |  |  |  |
|  | W132 | Strong VDW |  |  |  |
| F | I72 | Strong VDW | - | - | - |
| Y | L123 | Strong VDW | - | - | - |
|  | L80 | Strong VDW |  |  |  |
|  | N76 | H-bond |  |  |  |
|  | Y83 | H-bond |  |  |  |
|  | L94 | Strong VDW |  |  |  |
|  | S142 | H-bond |  |  |  |
| VDW: van der Waals (VDW) forces; H- bond: hydrogen bonds | | | | | |
