## Supplementary material for "Inferring MHC interacting SARS-CoV-2 epitopes recognized by TCRs towards designing T cell-based vaccines": Table S3

| **Table S3: Detailed interacting information of the hit peptide antigens of the N query of SARS-CoV-2 in models of TCR-pMHC complex for HLA-B3501** | | | | | |
| --- | --- | --- | --- | --- | --- |
| **Peptide  amino acid** | **MHC molecule** | | **TCR molecules** | | |
|  | **Amino acid** | **Bond type** | **Chain** | **Amino acid** | **Bond type** |
| N | R62 | H-bond | D | Y96 | H-bond |
|  | Y171 | H-bond |  |  |  |
|  | Y7 | H-bond |  |  |  |
|  | Y159 | H-bond |  |  |  |
| S | Y159 | Strong VDW | - | - | - |
|  | Y7 | Strong VDW |  |  |  |
|  | Y99 | Strong VDW |  |  |  |
| S | Y99 | H-bond | - | - | - |
|  | L156 | Strong VDW |  |  |  |
| P | - | - | D | Y96 | Strong VDW |
| D | Q155 | Strong VDW | - | - | - |
| D | - | - | - | - | - |
| Q | - | - | E | N28 | H-bond |
|  |  |  | E | R30 | H-bond |
| I | - | - | - | - | - |
| G | - | - | - | - | - |
| Y | W147 | H-bond | - | - | - |
| Y | S77 | H-bond | - | - | - |
|  | Y74 | Strong VDW |  |  |  |
|  | Y123 | Strong VDW |  |  |  |
|  | S116 | H-bond |  |  |  |
|  | W147 | Strong VDW |  |  |  |
| VDW: van der Waals (VDW) forces; H- bond: hydrogen bonds | | | | | |
