## Supplementary material for "Inferring MHC interacting SARS-CoV-2 epitopes recognized by TCRs towards designing T cell-based vaccines": Table S4

| **Table S4: Detailed interacting information of the hit peptide antigens of the S query of SARS-CoV-2 in models of TCR-pMHC complex for HLA-B3508** | | | | | |
| --- | --- | --- | --- | --- | --- |
| **Peptide  amino acid** | **MHC molecule** | | **TCR molecules** | | |
|  | **Amino acid** | **Bond type** | **Chain** | **Amino acid** | **Bond type** |
| L | W167 | Strong VDW | - | - | - |
|  | L163 | Strong VDW |  |  |  |
|  | Y159 | H-bond |  |  |  |
|  | Y7 | H-bond |  |  |  |
|  | Y171 | H-bond |  |  |  |
| Q | Y99 | Strong VDW | - | - | - |
|  | Y159 | Strong VDW |  |  |  |
|  | Y7 | Strong VDW |  |  |  |
| D | R156 | Strong VDW | - | - | - |
|  | R97 | H-bond |  |  |  |
|  | Y99 | H-bond |  |  |  |
| V | - | - | D | Y100 | Strong VDW |
| V | I66 | Strong VDW | D | Y100 | Strong VDW |
|  |  |  | D | F99 | Strong VDW |
| N | Q155 | Strong VDW | D | Y100 | Strong VDW |
|  |  |  | E | N32 | H-bond |
|  |  |  | D | F99 | Strong VDW |
| Q | - | - | D | Y100 | Strong VDW |
|  |  |  | E | N32 | Strong VDW |
|  |  |  | D | N101 | Strong VDW |
|  |  |  | E | Y35 | H-bond |
|  |  |  | D | Y35 | H-bond |
|  |  |  | D | S97 | H-bond |
| N | - | - | E | N32 | H-bond |
| A | - | - | E | N32 | Strong VDW |
|  |  |  | E | N30 | H-bond |
| Q | - | - | - | - | - |
| A | W147 | Strong VDW | - | - | - |
| L | W147 | H-bond | - | - | - |
| N | W147 | Strong VDW | - | - | - |
|  | S77 | H-bond |  |  |  |
|  | Y74 | H-bond |  |  |  |
|  | Y123 | Strong VDW |  |  |  |
|  | S116 | H-bond |  |  |  |
|  | K146 | H-bond |  |  |  |
| VDW: van der Waals (VDW) forces; H- bond: hydrogen bonds | | | | | |
