## Supplementary material for "Inferring MHC interacting SARS-CoV-2 epitopes recognized by TCRs towards designing T cell-based vaccines": Table S5

| **Table S5: Detailed interacting information of the hit peptide antigens of the S query of SARS-CoV-2 in models of TCR-pMHC complex for HLA-B4405** | | | | | |
| --- | --- | --- | --- | --- | --- |
| **Peptide  amino acid** | **MHC molecule** | | **TCR molecules** | | |
|  | **Amino acid** | **Bond type** | **Chain** | **Amino acid** | **Bond type** |
| N | R170 | H-bond | - | - | - |
|  | Y171 | H-bond |  |  |  |
|  | R62 | H-bond |  |  |  |
|  | Y159 | H-bond |  |  |  |
| K | Y9 | H-bond | - | - | - |
|  | K45 | H-bond |  |  |  |
|  | Y99 | H-bond |  |  |  |
|  | E63 | H-bond |  |  |  |
| K | Q155 | H-bond | - | - | - |
|  | Y159 | Strong VDW |  |  |  |
|  | Y99 | H-bond |  |  |  |
| F | I66 | Strong VDW | D | Y93 | Strong VDW |
| L | V152 | Strong VDW | D | Y31 | Strong VDW |
|  |  |  | D | W90 | Strong VDW |
|  |  |  | D | Y93 | Strong VDW |
| P | - | - | E | R94 | H-bond |
|  |  |  | E | R30 | H-bond |
| F | A150 | Strong VDW | D | Y31 | Strong VDW |
|  |  |  | E | Y100 | Strong VDW |
| Q | V152 | Strong VDW | - | - | - |
| Q | E76 W147 | H-bond H-bond | E | Y95 | Strong VDW |
|  |  |  | E | D98 | H-bond |
|  |  |  | E | D97 | H-bond |
| F | T143 | H-bond | - | - | - |
|  | Y123 | Strong VDW |  |  |  |
|  | Y116 | Strong VDW |  |  |  |
|  | Y84 | H-bond |  |  |  |
|  | I95 | Strong VDW |  |  |  |
| VDW: van der Waals (VDW) forces; H- bond: hydrogen bonds | | | | | |
