## Supplementary material for "Inferring MHC interacting SARS-CoV-2 epitopes recognized by TCRs towards designing T cell-based vaccines": Table S6

| **Table S6: Detailed interacting information of the hit peptide antigens of the S query of SARS-CoV-2 in models of TCR-pMHC complex for HLA-B0801** | | | | | |
| --- | --- | --- | --- | --- | --- |
| **Peptide  amino acid** | **MHC molecule** | | **TCR molecules** | | |
|  | **Amino acid** | **Bond type** | **Chain** | **Amino acid** | **Bond type** |
| F | Y59 | Strong VDW | - | - | - |
|  | W167 | Strong VDW |  |  |  |
|  | Y159 | H-bond |  |  |  |
|  | Y7 | H-bond |  |  |  |
|  | Y171 | H-bond |  |  |  |
|  | I66 | Strong VDW |  |  |  |
| S | I66 | Strong VDW | - | - | - |
|  | F36 | Strong VDW |  |  |  |
|  | N63 | H-bond |  |  |  |
| T | Y99 | H-bond | - | - | - |
|  | D156 | H-bond |  |  |  |
|  | Y159 | Strong VDW |  |  |  |
|  | N70 | H-bond |  |  |  |
| F | - | - | - | - | - |
| K | D9 | H-bond | - | - | - |
|  | T73 | H-bond |  |  |  |
|  | N70 | H-bond |  |  |  |
|  | D74 | H-bond |  |  |  |
| C | - | - | E | Q96 | H-bond |
|  |  |  | D | Y96 | Strong VDW |
| Y | W147 | Strong VDW | E | A97 | Strong VDW |
|  |  |  | D | H47 | Strong VDW |
|  |  |  | D | H32 | Strong VDW |
|  |  |  | D | L90 | Strong VDW |
| G | W147 | H-bond | E | Y98 | H-bond |
| V | S77 | H-bond | - | - | - |
|  | Y84 | H-bond |  |  |  |
|  | L81 | Strong VDW |  |  |  |
|  | W147 | Strong VDW |  |  |  |
|  | T143 | H-bond |  |  |  |
|  | L95 | Strong VDW |  |  |  |
| VDW: van der Waals (VDW) forces; H- bond: hydrogen bonds | | | | | |
