## Supplementary material for "Inferring MHC interacting SARS-CoV-2 epitopes recognized by TCRs towards designing T cell-based vaccines": Table S8

| **Table S8: Summary of the interatomic interactions within chain C and their binding site interactions in the TCR-pMHC complex of bat-CoV proteins** | | | | | | | |
| --- | --- | --- | --- | --- | --- | --- | --- |
| **Interatomic interactions** | | **HLA-A0201** | **HLA-B0801** | **HLA-B3501** | **HLA-B3508** | **HLA-B4405** | **HLA-E** |
|  |  | **N** | **S** | **N** | **S** | **S** | **N** |
| Mutually Exclusive Interactions | VDW interactions | 17 | 9 | 14 | 16 | 14 | 34 |
|  | VDW clash interactions | 100 | 64 | 67 | 56 | 52 | 130 |
|  | Covalent interactions | 7 | 3 | 2 | 1 | 3 | 6 |
|  | Covalent clash interactions | 1 | 1 | 0 | 0 | 0 | 2 |
|  | Proximal | 878 | 640 | 702 | 926 | 776 | 878 |
|  | Total | 1003 | 717 | 785 | 999 | 845 | 1059 |
| Polar Contacts | Polar contacts | 16 | 20 | 16 | 23 | 17 | 23 |
|  | Weak polar contacts | 31 | 23 | 13 | 26 | 19 | 43 |
|  | Total | 47 | 43 | 29 | 49 | 36 | 66 |
| Feature Contacts | Hydrogen bonds | 15 | 16 | 10 | 18 | 13 | 15 |
|  | Weak hydrogen bonds | 13 | 9 | 8 | 10 | 5 | 14 |
|  | Ionic interactions | 0 | 0 | 0 | 0 | 1 | 3 |
|  | Aromatic contacts | 40 | 4 | 0 | 0 | 7 | 99 |
|  | Hydrophobic contacts | 99 | 86 | 49 | 62 | 74 | 151 |
|  | Carbonyl interactions | 1 | 0 | 1 | 2 | 0 | 0 |
|  | Total | 168 | 115 | 68 | 92 | 100 | 282 |
| Chain C: Hit peptide; VDW: van der Waals (VDW) forces; N: Nucleocapsid phosphoprotein; S: Surface glycoprotein; HLA: Human leukocyte antigen | | | | | | | |
