## Supplementary material for "Inferring MHC interacting SARS-CoV-2 epitopes recognized by TCRs towards designing T cell-based vaccines": Table S9

| **Table S9: Summary of the interatomic interactions within chain C and their binding site interactions in the TCR-pMHC complex of MERS-CoV proteins** | | | | | | | |
| --- | --- | --- | --- | --- | --- | --- | --- |
| **Interatomic interactions** | | **HLA-A0201** | **HLA-B0801** | **HLA-B3501** | **HLA-B3508** | **HLA-B4405** | **HLA-E** |
|  |  | **N** | **S** | **N** | **S** | **S** | **N** |
| Mutually Exclusive Interactions | VDW interactions | 22 | 11 | - | - | 14 | 32 |
|  | VDW clash interactions | 92 | 35 | - | - | 36 | 126 |
|  | Covalent interactions | 2 | 1 | - | - | 1 | 6 |
|  | Covalent clash interactions | 0 | 0 | - | - | 0 | 2 |
|  | Proximal | 843 | 690 | - | - | 731 | 844 |
|  | Total | 959 | 737 | - | - | 782 | 1010 |
| Polar Contacts | Polar contacts | 14 | 19 | - | - | 19 | 21 |
|  | Weak polar contacts | 25 | 24 | - | - | 15 | 39 |
|  | Total | 39 | 43 | - | - | 34 | 60 |
| Feature Contacts | Hydrogen bonds | 13 | 17 | - | - | 14 | 14 |
|  | Weak hydrogen bonds | 10 | 7 | - | - | 6 | 14 |
|  | Ionic interactions | 0 | 5 | - | - | 9 | 0 |
|  | Aromatic contacts | 37 | 3 | - | - | 6 | 99 |
|  | Hydrophobic contacts | 72 | 52 | - | - | 74 | 153 |
|  | Carbonyl interactions | 1 | 0 | - | - | 0 | 0 |
|  | Total | 133 | 84 | - | - | 109 | 280 |
| Chain C: Hit peptide; VDW: van der Waals (VDW) forces; N: Nucleocapsid phosphoprotein; S: Surface glycoprotein; HLA: Human leukocyte antigen | | | | | | | |
